# Aryl N-acetamide compounds exert antimalarial activity by acting as agonists of rhomboid protease *Pf*ROM8 and cation channel *Pf*CSC1

**DOI:** 10.1101/2025.03.18.644039

**Authors:** Coralie Boulet, Joyanta K. Modak, Natalie A. Counihan, Molly Parkyn Schneider, William Nguyen, Madeline G. Dans, Claudia B.G. Barnes, Zahra Razook, Kirsty McCann, Alyssa E. Barry, Brendan S. Crabb, Brad E. Sleebs, Tania F. de Koning-Ward, Paul R. Gilson

## Abstract

With resistance to current frontline antimalarials spreading globally, new drug candidates need to be discovered to populate the antimalarial drug development pipeline. We previously screened the Medicines for Malaria Venture Pathogen Box for compounds that prevent *Plasmodium falciparum* parasites from exiting and invading human erythrocytes, steps essential for the proliferation of parasites in the blood, which causes disease. Compound MMV020512 (M-512) was identified in this screen and live cell imaging here established that it does not specifically inhibit invasion but likely inhibits intraerythrocytic parasite growth. M-512 resistance selection in parasites led to the identification of mutations in the membrane protease *Pf*ROM8 and the cation ion channel *Pf*CSC1. *Pf*ROM8 was validated as a target of M-512 when a L562R putative resistance mutation was engineered into wildtype parasites reproducing the resistance phenotype. Knockdown of wildtype *Pf*ROM8, the L562R mutant and CSC1 reduced parasite growth, indicating the proteins are functionally important. Counterintuitively, the *Pf*ROM8 and *Pf*CSC1 knockdown parasites became more resistant to M-512 suggesting that the compound is an agonist of both proteins which may form a functional complex and that dysregulation of this complex is deleterious to parasite growth.

## Introduction

Malaria is caused by infection with *Plasmodium* parasites and resulted in 263 million cases and 597,000 deaths in 2023 (World Health Organization 2023). The only means of clearing parasite infections is with antimalarial medicines consisting of small molecule inhibitory drugs. Artemisinin combination therapies (ACTs) are the frontline treatments for malaria, comprising a fast-acting but short-lived artemisinin derivative in combination with a slower-acting but longer-lasting partner drug (World Health Organization 2023). Treatment with ACTs is failing due to parasite resistance to both the artemisinin-based component and the partner drug, with resistance now widespread in South East Asia, Papua New Guinea and more recently detected in African regions (Dondorp, Nosten et al. 2009, Lautu-Gumal, Razook et al. 2021, Yoshida, Yamauchi et al. 2021, Tumwebaze, Conrad et al. 2022, Rosenthal, Asua et al. 2024). For this reason, there is a continued need to discover new drug targets and develop novel medicines against these targets that parasites are not already resistant to.

To facilitate this, large drug libraries have been screened for antimalarial activity with tens of thousands of compounds now known to act against the asexual blood stage of *P. falciparum,* the species responsible for most malaria deaths *(Gamo, Sanz et al. 2010, Guiguemde, Shelat et al. 2010)*. Medicines for Malaria Venture (MMV) has provided libraries of antimalarial compounds to researchers to assist in the identification of new drug targets and their mechanisms of action (Spangenberg, Burrows et al. 2013, Van Voorhis, Adams et al. 2016). This approach has helped identify many new drugs targets, but the process is slow and resource intensive, with many compounds acting against the same proteins, e.g. *Pf*ATP4 (Lehane, Ridgway et al. 2014, Dennis, Rosling et al. 2018). To help identify new targets that can also be used as tools to understand parasite biology, we have developed screens to rapidly identify inhibitors that block parasite egress from the host erythrocyte or invasion into another erythrocyte (Dans, Weiss et al. 2020). One advantage of invasion inhibitors is that they do not need to directly kill the parasites. Instead, they only need to prevent the parasites from successfully invading erythrocytes to stop parasite proliferation as the extracellular invasive merozoites lose viability very quickly and are highly susceptible to host immune responses (Bouharoun-Tayoun, Oeuvray et al. 1995, Boyle, Reiling et al. 2015, Arora, Hart et al. 2018).

To identify egress or invasion inhibitory compounds, we screened the 400-compound MMV Pathogen Box against *P. falciparum* over a four-hour period when schizont-stage parasites are in the process of rupturing to release merozoites that invade new erythrocytes. This screen identified a dozen merozoite egress inhibitors and about two dozen invasion inhibitors (Dans, Weiss et al. 2020). Once inhibitors with known or suspected targets were removed, several novel compounds were further investigated. This included MMV020291, which was shown to potently inhibit invasion (Dans, Weiss et al. 2020) by targeting actin-1 (PF3D7_1246200) and profilin (PF3D7_0932200) (Dans, Piirainen et al. 2023). MMV020291 appears to inhibit the polymerisation of actin, which is required by the merozoite to mechanically force its entry into erythrocyte (Dans, Piirainen et al. 2023). Another inhibitor, MMV006833, did not directly block invasion but instead was shown to affect the subsequent differentiation of newly invaded merozoites into larger intraerythrocytic amoeboid forms. The target of MMV006833 is the *Pf*START1 lipid transfer protein (also called PfPV6, PF3D7_0104200) (van Ooij, Withers-Martinez et al. 2013, Hill, Ringel et al. 2016, Fréville, Ressurreição et al. 2023, Dans, Boulet et al. 2024) indicating the inhibitor probably blocks *Pf*START1 from transporting lipids between membranes after invasion and thereby preventing the parasite from expanding and growing (Dans, Boulet et al. 2024).

Here we sought to understand the mechanism of action of MMV020512, another invasion inhibitor that also had modest activity against ring-stage and older trophozoite-stage parasites, suggesting the compound’s target might be expressed throughout the asexual cell cycle (Dans, Weiss et al. 2020). Selection for resistance in wild-type *P. falciparum* was successful, and mutations were observed in the rhomboid protease 8 (ROM8, PF3D7_1411200) and the calcium stress response 1 (CSC1, PF3D7_1250200) protein genes. Highly potent MMV020512 analogues were developed that appear to act as agonists, possibly by preventing closure of the channels and thus the dysregulation of the flow of cations. This was evident when both *Pf*ROM8 or *Pf*CSC1 were separately knocked down and rather than being sensitised to MMV020512, parasites became more resistant to the inhibitors. Although it was not established how MMV020512 analogues specifically blocked invasion, *Pf*CSC1 and *Pf*ROM8 may be important for parasite osmoregulation and this process may be required during merozoite development and invasion.

## Results

### M-512 does not appear to be a direct inhibitor of invasion

To determine if M-512 was acting as a direct inhibitor of erythrocyte invasion, live cell imaging of invading merozoites was performed in the presence of 10 x EC_50_ (∼5 µM) of M-512 (Figure S1A). Late-stage *P. falciparum* W2mef strain schizonts were placed in an environmental chamber and imaged as the merozoites exited and attempted to invade neighbouring uninfected erythrocytes. Fourteen schizont egresses were observed for M-512, as well as ten for the 0.5% DMSO control (Supplementary videos 1 and 2). Although there was a reduction in the average number of invasions, and productive invasions in which transformation into ring-stage parasites was observed following M-512 treatment, this was not significant (Figure S1B, p>0.05). The times for preinvasion (the time from first contact to the start of merozoite penetration), and the duration of penetration were also measured and were not significantly different between M-512 and the DMSO control (Figure S1B, p>0.05). With live cell imaging unable to provide insights into the mechanism of action of M-512 we decided to select for resistance to the compound and find mutations in the possible target protein.

### Selection of parasites that are resistant to M-512

We attempted to select for resistance in five populations (A-E), each comprising 1x 10^8^ asexual blood stage 3D7 wild-type parasites. Briefly the parasites were treated with 5 µM M-512 until they began to die, at which point M-512 treatment was halted and the population allowed to recover. This was repeated for a total of three times with the parasites appearing to recover faster each round. These parasite populations were subjected to a 72-hour growth assay in serially diluted M-512 compound with lactate dehydrogenase (LDH) activity measured to quantify growth. The increase in resistance, as indicated by growth EC_50_, ranged from 6- to 100-fold greater than the parental 3D7 parasites (EC_50_ 0.39 µM) with populations D, A and E (PopD, -A and -E) the most resistant in that order (Figure S2A,B). PopA also exhibited a unusual phenomenon where its growth appeared to improve at intermediate M-512 concentrations (0.04 – 2.5 µM) before falling at higher compound concentrations (5 - 10 µM) (Figure S2A,B). For these reasons we chose PopA, PopD and PopE for the selection of clonal lines for further analysis.

Two clonal lines isolated from each of the three populations were subjected to further 72-hour growth assays. The clonal parasites all retained their resistance indicating the resistance was stable and heritable (Figure 1A). Furthermore, PopA and PopE cloned lines still exhibited an increase in growth above the basal rate in intermediate concentrations of M-512 before becoming greatly inhibited at higher concentrations.

**Figure 1.**
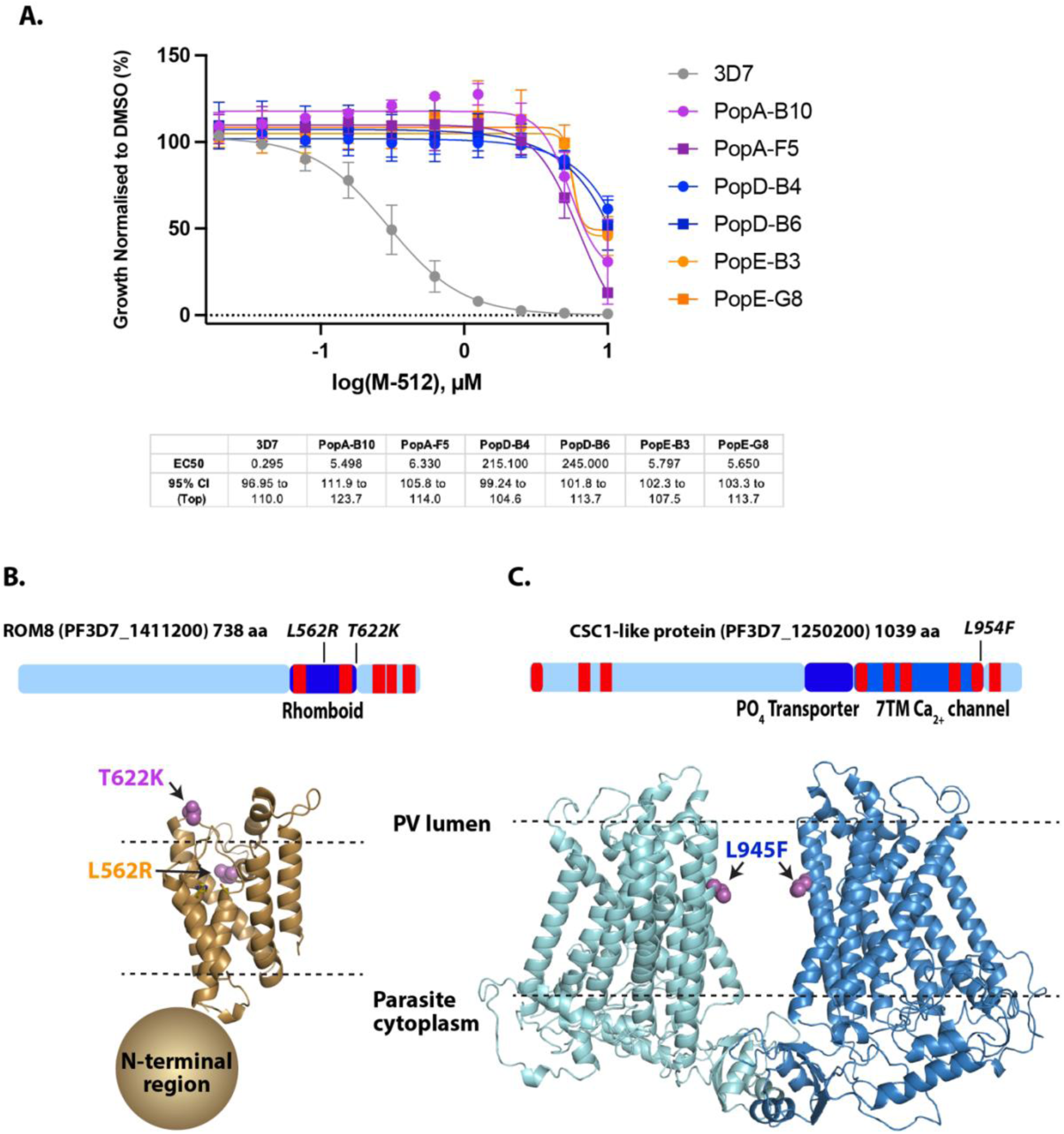
M-512 resistant parasites had mutations in*Pf*ROM8 and *Pf*CSC1. **A)** Growth curves of clones of resistant parasites treated with a dilution series of M-512 for 72 hours. Growth was assessed by measuring lactate dehydrogenase activity of the parasite infected erythrocytes. EC_50_ values (µM) and 95% confidence intervals are shown below the graph. The data represent the average of three biological replicates in technical triplicate with error bars indicating SD. Note that the EC_50_ values for PopD and PopE may be underestimated as the inhibition curves do not fall below 50% at the highest M-512 concentration tested (10 µM). **B, Top)** Diagram of *Pf*ROM8 showing position of putative M-512 resistance mutations, predicted functional domains (dark blue) and transmembrane regions (red). **B, Bottom)** Alpha-fold generated structure of the rhomboid protease domain of *Pf*ROM8. The structure of the N-terminal region was not confidentially predicted and is represented as a sphere. **C, Top)**. As per panel B but for *Pf*CSC1 protein. **C, Bottom)** Alpha-fold generated structure of a predicted *Pf*CSC1 dimer showing locations of M-512 resistance mutations.

### Whole genome sequencing revealed potential resistance mutations in three proteins

Genomic DNA from the two clonal lines of each population as well as the 3D7 parental control, was extracted and subjected to whole genome sequencing using MinION technology (Dans, Piirainen et al. 2023, Harrison, Mehra et al. 2024). Non-synonymous single nucleotide polymorphisms (SNPs) were identified within the coding sequencing of several genes that are listed in Table S1. The mutations that were shared in both clonal lines from each population were considered most likely to be responsible for M-512 resistance. In PopA, both clones had a T622K mutation in rhomboid protease 8 (*Pf*ROM8, PF3D7_1411200) resulting from a ACG to AAG change in a threonine codon (Figure 1B and Table S1). PopE clones also had a *Pf*ROM8 mutation but in a different position at L562R, in which the leucine codon CTC was changed to the arginine codon CGC (Figure 1B and Table S1). Both clones of PopD harboured a L954F mutation in the calcium permeable stress-gated cation channel 1-like protein *Pf*CSC1 (*Pf*CSC1, PF3D7_1250200) resulting from a TTG to TTT change in a leucine codon (Figure1C and Table S1). The PopD parasites also had a M1061I mutation in the valine tRNA ligase gene (Val tRNA Lig, PF3D7_1461900) from an ATG to ATA mutation in the methionine codon (Table S1).

To validate the presence of these mutations, short regions of DNA flanking the mutations were amplified by PCR and sequenced (Table S2). This revealed that all mutations were present as per the whole genome sequencing (Table S1 and Figure S3). *Pf*ROM8 is essential (Ramaprasad and Blackman 2024) and is a rhomboid-type multi-pass transmembrane serine protease that acts upon the transmembrane regions of other proteins (Figure 1B). The *P. falciparum* genome encodes eight rhomboid proteases with *Pf*ROM1 and *Pf*ROM4 predicted to help remove the merozoite surface coat during erythrocyte invasion (Howell, Hackett et al. 2005, Baker, Wijetilaka et al. 2006). *Pf*ROM8, however, appears to be involved in merozoite development which could in turn affect their ability to invade (Ramaprasad and Blackman 2024). *Pf*CSC1 is predicted to be an essential multi-pass transmembrane channel specific for cations (Zhang, Wang et al. 2018) (Figure 1C). It belongs to a family of seven pass transmembrane domain channels that may have mechanosensitive or osmoregulatory roles (Hou, Tian et al. 2014). A specific role for Val-tRNA Ligase in invasion is difficult to envisage since such an enzyme would be expected to have a predominately housekeeping role. It is possible that Val tRNA Ligase may also be targeted by M-512 or that it compensates for the resistance mutations in *Pf*CSC1. As we were interested proteins with a putative invasion role, we focussed our investigation into the *Pf*ROM8 and *Pf*CSC1 mutations for the remainder of the study.

### Mutations in PfROM8 are associated with changes in parasite fitness

To determine if the M-512 resistance mutations were associated with a change of fitness, LDH activity was measured at the trophozoite stage (LDH_0h_) and 48 hours later when the parasite were trophozoites again (LDH_48h_). Parasite growth was expressed as the ratio of LDH_48h_/LDH_0h_ and this was repeated for up to four consecutive cell cycles in four independent experiments (Figure 2). The results indicated that, in the absence of M-512, clonal line PopD-B4 (*Pf*CSC1 L954F) did not grow at a significantly different rate than the 3D7 parental parasites (Figure 2). PopA-B10 (*Pf*ROM8 L562R) and PopE-B3 (*Pf*ROM8 L562R) appeared to grow more slowly than the other parasite lines however this only reached significance for PopE-B3 (*Pf*ROM8 L562R).

**Figure 2.**
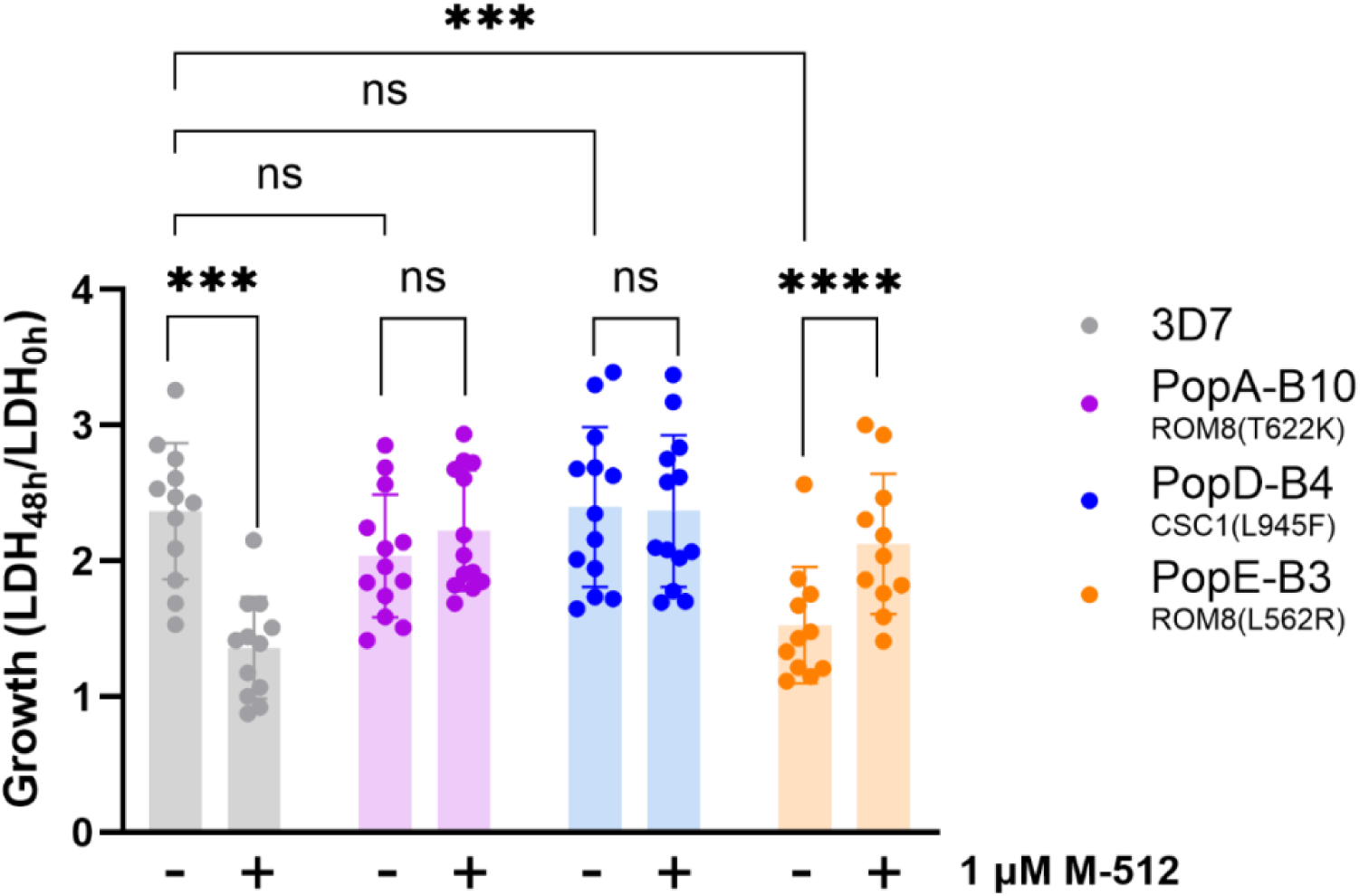
M-512 resistance mutations can improve the growth of parasites in the presence of M-512. The growth rates of wild-type (3D7) and M-512 resistant clonal parasite lines were derived by measuring LDH activity at the trophozoite stage (LDH_0h_) and again 48 hours later at the following trophozoite stage (LDH_48h_). The fold increase in LDH levels after each 48-hour cycle is shown -/+ 1 µM M-512, with bar height indicating the mean of individual LDH48h/LDH0h values and errors bars indicating the SD. The data were obtained from four independent experiments each comprising 2 - 4 consecutive cycles with each data point representing a separate 48-hour cycle. Statistical analysis between parasite lines was performed in PRISM V10 using Ordinary one-way ANOVA with Dunnett’s multiple comparisons test, and for growth between each line -/+ M-512, paired t tests were used. *p<0.05,**p<0.01, **p<0.001 and ****p<0.0001. ns, not significant.

In parallel, parasite growth rates were measured in the continued presence of 1 µM M-512 (equivalent to 2-3 x EC_50_) and, as anticipated, the sensitive 3D7 parasites grew poorly in the presence of M-512 (Figure 2). The PopD-B4 (*Pf*CSC1 L954F) mutant lines grew equally well +/- M-512 due to their resistance (p>0.05). In contrast, PopA-B10 (*Pf*ROM8 T622K) and PopE-B3 (*Pf*ROM8 L562R) mutants grew better in the presence of M-512 but this was only significant for PopE-B3 (Figure 2, p<0.05)). This indicates that low or intermediate concentrations M-512 could potentially act as an agonist of the less functional mutant *Pf*ROM8 proteins, increasing their activity and partly restoring parasite growth.

### Introduction of the PfROM8 L562R SNP into 3D7 parasites increased resistance to M-512

To validate that *Pf*ROM8 L562R or T622K mutations were responsible for resistance to M-512, we attempted to introduce these into the *Pf*ROM8 locus of wild-type parasites and determine if M-512 resistance was conferred. The mutations, as well as a *Pf*ROM8(WT) control, were each introduced separately into a synthetic recodonised block covering the last 648 bp of *Pf*ROM8’s coding sequence from which the introns had been removed (Figure S4A). This was sewn in frame to a 941 bp upstream block identical to the native gene which together formed the whole 5’ homology block that was inserted upstream of the human dihydrofolate reductase (hDHFR) selection marker in plasmid p1.2 (Figure S4A) (Marapana, Dagley et al. 2018, Dans, Piirainen et al. 2023). A 3’ homology block derived from the 3’ region of *Pf*ROM8’s coding sequence was cloned downstream of the hDHFR cassette and the resulting donor construct was transfected into parasites, along with recombinant Cas9 and a synthetic guide RNA specific for *Pf*ROM8 (McHugh, Bulloch et al. 2023). Under selection with the antifolate WR99210, the donor plasmid was designed to replace the *Pf*ROM8 locus and introduce the mutations by double cross-over recombination (Figure S4A). Using this approach, we recovered transfected parasites CR-ROM8(WT), CR-ROM8(L562R) and CR-ROM8(T622K). To ensure that the *Pf*ROM8 locus had been altered as intended, diagnostic PCRs were performed with primers specific for the introduced sequence flanking the mutations. These produced bands of the expected size for CR-ROM8(WT), CR-ROM8(L562R) and CR-ROM8(T622K) parasites, whereas no product was obtained for 3D7 as expected (Figure S4B, top). Sequencing of these products revealed they contained the desired mutations (Figure S4C). Longer amplification products obtained by PCR using primers specific for the native flanking sequences outside the homology blocks produced correctly sized products for CR-ROM8(WT) and CR-ROM8(L562R) but not for CR-ROM8(T622K) and the parental 3D7 (Figure S4B, middle and bottom). Sequencing of the CR-ROM8(WT) and CR-ROM8(L562R) PCR products matched that expected for correct integration into the *pfrom8* locus while the CR-ROM8(T622K) product had not integrated into *pfrom8* but into the gene of a conserved membrane protein of unknown function (PF3D7_0719900). We repeated the *Pf*ROM8 T622K transfection again with a different guide RNA but without success and so proceeded to just analyse the correctly modified CR-ROM8(WT) and CR-ROM8(L562R) parasites.

To determine if the CRISPR/Cas9 engineered *Pf*ROM8 mutation conferred resistance to M-512, a 72-hour growth assay was performed on parasites challenged with a dilution series of the inhibitor (Figure S5A). As anticipated, the EC_50_ of CR-ROM8(WT) parasites was like those of the 3D7 parental line and the incorrectly modified CR-ROM8(T622K) parasites. This supported the PCR and DNA sequencing data which indicated that the *pfrom8* locus in the CR-ROM8(T622K) parasites had not been modified and no further work was done with these parasites.

Next, clonally derived CR-ROM8(WT) and CR-ROM(L562R) parasites were subjected to 72-hour growth assays. Once again, the CR-ROM8(L562R) parasites were >15 fold more resistant to M-512 than CR-ROM8(WT) control line (Figure 3A). We also noticed that the growth inhibition of the CR-ROM8(L562R) parasites at mid-range concentrations of M-512 (0.31 – 5 µM) increased above the 100% DMSO control for normalised data (Figure 3A) indicating a possible improvement in the growth rate.

**Figure 3.**
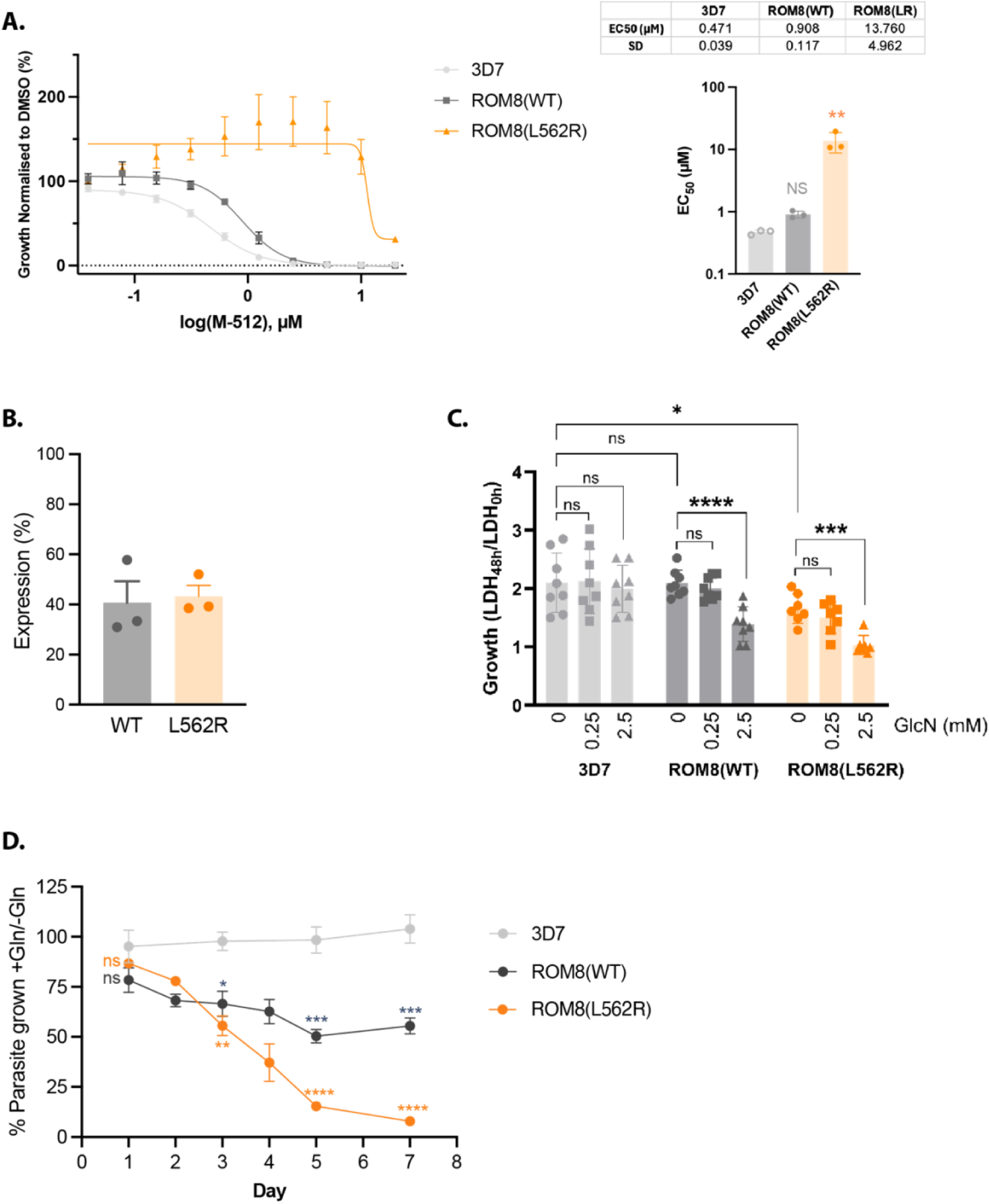
Knockdown of *Pf*ROM8 reduces parasite growth. A, Left) 72-hour growth inhibition assay of parasites challenged with M-512 indicated CR-ROM8(L562R) parasites were more resistant than the control CR-ROM8(WT) parasites and wildtype 3D7 parasites. Growth curves fitted using linear regression in PRISM V10. **Right)** LDH activity was used to quantify parasite growth and the EC50 of the CR-ROM8(L562R) parasites was significantly greater than the CR-ROM8(WT) and 3D7 parasites. The data represent three biological replicates each of technical duplicates. **B)** QPCR of *rom8* gene in CR-ROM8(WT) and CR-ROM8(L562R) parasites treated +/ 2.5 mM GlcN showing GlcN knocks down mRNA levels to about 40% levels of untreated parasites. 2.5 mM GlcN was added to ring-stage parasites and mRNA was harvested at the schizont-stage of the same cycle. The data represent 3 biological replicates. **C)** 2.5 mM and 0.25 mM glucosamine (GlcN) were added to trophozoite stage 3D7, CR-ROM8(WT) and CR-ROM8(L562R) parasites (0h) which were harvested 48 hours later (48h). LDH activity was measured and the ratio of LDH_48h_/LDH_0h_ is shown for each cycle. The experiment was performed three times with consecutive cycles in each experiment and each data point corresponds to the LDH_48h_/LDH_0h_ ratiofor a different 48h interval. Bar height indicates mean, and whiskers show SD. Comparison of GlcN treatment within lines were tested with RM one-way ANOVA with Dunnett’s multiple comparisons test. Comparison of growth in absence of GlcN (0 mM) was tested with an ordinary one-way ANOVA with Dunnett’s multiple comparisons test. **D)** Multicycle growth assay of 3D7, CR-ROM8(WT) and CR-ROM8(L562R) parasites following treatment +/- 2.5 mM GlcN. Each day growth was estimated by SYBR green fluorescence with the data points representing the fluorescence ratio of +GlcN/-GlcN. Comparison of growth ratios between 3D7 and the transgenic lines was tested with an ordinary one-way ANOVA with Dunnett’s multiple comparisons test. Tests performed in PRISM V10 with *p<0.05, **p<0.01, **p<0.001 and ****p<0.0001. ns, not significant.

### Knockdown of PfROM8-glmS reduced parasite growth

The modification of the *Pf*ROM8 locus also introduced a *glmS* riboswitch to enable the knockdown of *Pf*ROM8 expression following addition of glucosamine (GlcN), which activates *glmS* to cleave its mRNA, thereby reducing protein expression (Prommana, Uthaipibull et al. 2013). We have previously observed that reduction of target protein levels can sensitise the parasites to the effects of the inhibitor, helping to confirm the inhibitor’s mechanism of action (Dans, Boulet et al. 2024). Before conducting this experiment, we attempted to measure the knockdown level of *Pf*ROM8 following treatment with 2.5 mM GlcN by detecting a single hemagglutinin (HA) epitope tag appended to the C-terminal end of *Pf*ROM8. However, western blot analysis and immunofluorescence microscopy experiments with HA antibodies failed to detect signal (Figure S5B,C). Careful examination of the HA epitope coding sequence revealed the last amino acid (alanine) was missing perhaps accounting for why the monoclonal mouse antibody would not recognise it. We then constructed a new donor plasmid in which we attempted to append three copies of the HA epitope to *Pf*ROM8, but transfected parasites could not be obtained despite repeated attempts.

Without an epitope tag on ROM8 with which to quantify its knockdown, we used qPCR with primers specific to the gene to quantify its down regulation. 2.5 mM glucosamine (GlcN) was added to ring-stage CR-ROM8(WT) and CR-ROM8(L562R) parasites and they were grown until the schizont-stage followed by harvesting of their RNA. GlcN activates the *glmS* ribozyme causing *in cis* cleavage of *rom8’s* 3’UTR and destabilising its mRNA resulting in lower protein expression (Prommana, Uthaipibull et al. 2013). As a control, CR-ROM8 parasites untreated with GlcN were grown in parallel and the mRNA levels +GlcN/-GlcN indicated that 2.5 mM GlcN treatment caused *pfrom8s’* mRNAs to decrease to about 40% of the untreated parasites for both WT and L652R lines (Figure 3B). We do not know what degree of protein knockdown this reduction in mRNA equates to but for other genes we have previously tagged, the protein knockdown was to about 20% the levels of parasites untreated with GlcN (Dans, Piirainen et al. 2023, Dans, Boulet et al. 2024).

To measure the growth rates of the engineered parasite lines the LDH activities were again assessed as per Figure 2. As anticipated, the CR-ROM8(WT) parasites grew similarly to the parental 3D7 parasites with the CR-ROM8(L562R) parasites growing at a reduced rate (Figure 3C). Addition of 2.5 mM GlcN caused a significant reduction in growth for both CR-ROM(WT) and CR-ROM8(L562R) parasites, with 0.25 mM GlcN having an intermediate effect probably due to less knockdown (Figure 3C). These parasites were also subjected to a multicycle assay treated +/- 2.5 mM GlcN and growth was estimated by SYBR green fluorescence. Wildtype 3D7 parasites that were not tagged with *glmS* were also included and the ratio of +GlcN/-GlcN values indicates that growth of both the CR-ROM8 parasites significantly decreased following GlcN treatment compared to 3D7 which was unaffected (Figure 3D). Giemsa smears from the multicycle assay were also examined and indicate that the CR-ROM8(WT) parasites slowed by Day 3 at the trophozoite stage but appeared to keep growing whereas the CR-ROM8(L562R) parasites appeared to stop growing at trophozoites on Day 3 (Figure S6).

### Knockdown of PfROM8 improves resistance to M-512 rather than sensitising the parasites

Next, we examined the effects of *Pf*ROM8 knockdown with GlcN in combination with M-512 treatment. After growth for 72 hours the raw LDH activity values were plotted rather than normalised data so as not to mask the effects of the treatments. For 3D7 parasites the addition of GIcN did not alter their EC_50_s suggesting a negligible effect on parasite growth (Figure 4A). At low concentrations of M-512 (0.04 – 0.31 µM), CR-ROM8(WT) parasites grew more poorly in 2.5 mM GlcN than in 0 mM or 0.25 mM GlcN. In contrast, at higher concentrations of M-512 (0.63 - 2.5 µM), the CR-ROM8(WT) parasites grew better in GlcN than in the absence of GlcN. This resulted in higher EC50s for CR-ROM8(WT) in 0.25 mM or 2.5 mM GlcN than in 0 mM GlcN. The effect of 0.25 mM GlcN treatment was between that of 0 mM and 2.5 mM GlcN (Figures 4A and S7A,C). This suggests that in GlcN-treated CR-ROM8(WT) parasites, which have reduced levels of *Pf*ROM8, M-512 stimulation of *Pf*ROM8 activity is not as detrimental to parasite growth as it is in non-GlcN treated parasites which have much higher levels of *Pf*ROM8 (Figure 4A and S7A,C). M-512 is therefore likely acting as an agonist of *Pf*ROM8 activity which, if too strongly stimulated, tends to inhibit parasite growth.

**Figure 4.**
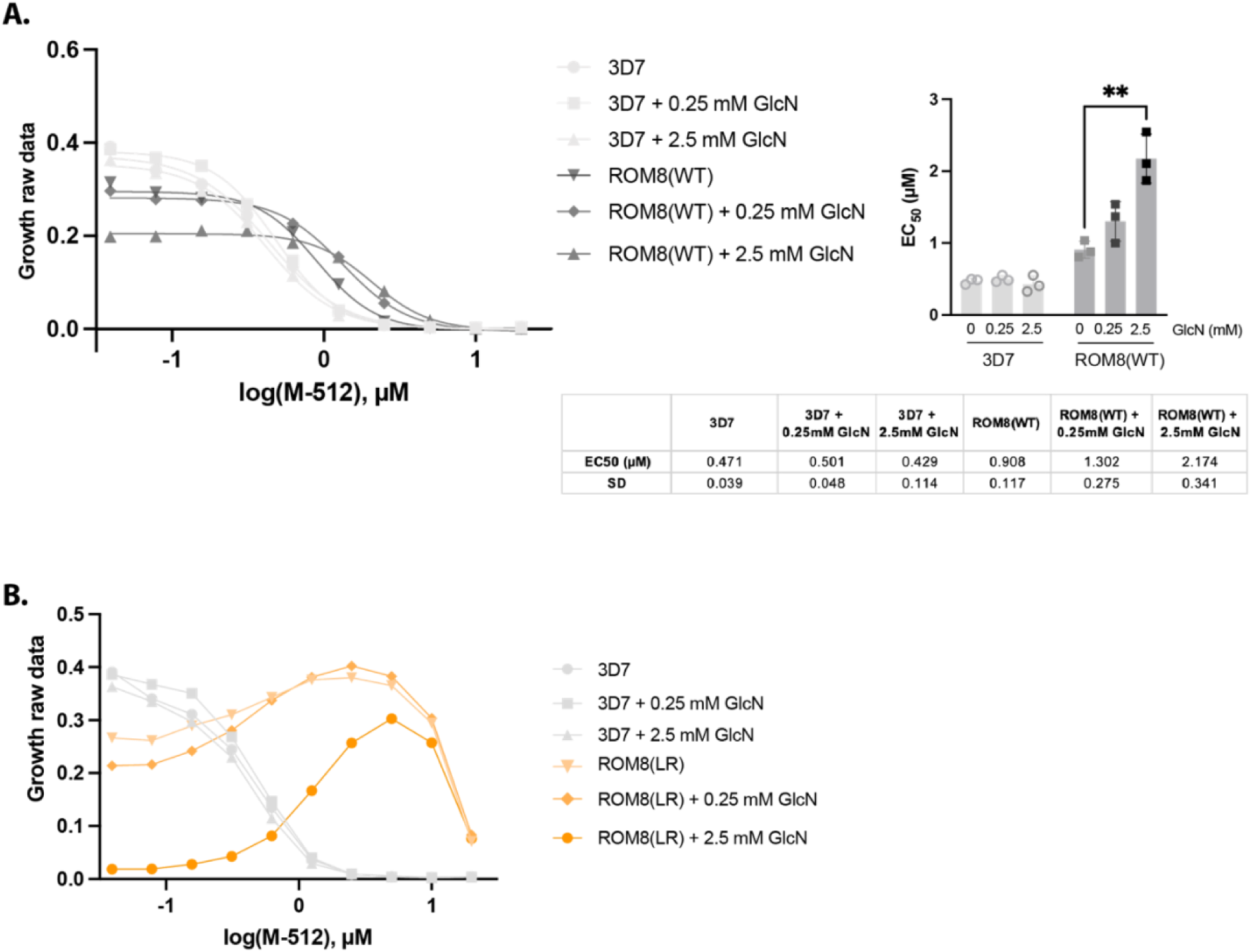
Introduction of a L562R mutation into *Pf*ROM8 and knocking down its expression reduced parasite growth which could be partly reversed though supplementation with M-512. A, Left) Growth inhibition curves showing raw LDH activity (OD 650 nm) with parasites treated with 0.25 and 2.5 mM GlcN which knocked down CR-ROM8(WT) expression. Error bars indicating SD values have been omitted for clarity but a complete version with SD whiskers is shown in Figure S7A. **Right)** GlcN had little effect upon the M-512 EC_50_s of 3D7 parasites but knockdown of CR-ROM8(WT) increased the EC_50_ of M-512 to being almost two-fold greater in 2.5 mM GlcN than 0 mM GlcN. **B)** Examination of the raw LDH values versus M-512 indicated that growth of CR-ROM8(L562R) parasites was reduced by knockdown with 2.5 mM GlcN and this was partly reversed by addition of intermediate concentrations of M-512 (2.5–10 µM). Note it was not possible to fit linear regression curves to this data (and to derive growth EC_50_s) and so connecting lines are instead shown between data points. A complete version of this data showing SD whiskers is shown in Figure S7B. Growth curves represent three biological replicates of technical duplicates and in **A)** one-way ANOVA with a Dunnett’s multiple comparison test was used. **p<0.01.

For CR-ROM8(L562R) parasites the effects of *Pf*ROM8 knockdown and M-512 levels were a more extreme version of that observed for CR-ROM8(WT), preventing us from calculating their EC_50_s (Figure 4B and S7B,C). At low M-512 concentrations (0.04 – 0.63 µM) and 2.5 mM GlcN, the CR-ROM8(L562R) parasites grew very poorly, possibly because the mutant *Pf*ROM8 protein was both less functional (due to the mutation) and was expressed at reduced levels (Figure 4C, Figure S7B,C). At intermediate concentrations of M-512 (1.25 – 5 µM), the growth of CR-ROM8(L562R) parasites improved, possibly because the compound increased the functionality of the residual levels of *Pf*ROM8(L562R) protein. At low concentrations of M-512, treatment of *Pf*ROM8(L562R) parasites with 0.25 mM GlcN only slightly reduced growth relative to 0 mM GlcN parasites probably because *Pf*ROM8 was knocked down less than 50% (Elsworth, Matthews et al. 2014). As the concentration of M-512 increased from 0.04-2.5 µM, the growth of CR-ROM8(L562R) parasites improved, both in the presence and absence of GlcN, suggesting that the putative *Pf*ROM8 agonist was improving the function of the mutant protein. At high M-512 concentrations (5 and 20 µM) the growth of all CR-ROM8(L562R) parasites was supressed as the agonistic effects of M-512 possibly became too strong.

### Knockdown of PfCSC1 reduces parasite growth

To determine if PfCSC1 was essential also important for growth, reverse genetics was employed to deplete CSC1 expression using the riboswitch system. Here, the *pfcsc1* locus was targeted by transfecting *P. falciparum* 3D7 with the pCSC1-HAglmS construct in conjunction with Cas9 protein and a synthetic guide RNA (Figure S8A). Positive transfectants were cloned by limiting dilution to ensure a pure population of CR-CSC1(WT) parasites. Diagnostic PCR confirmed that clonal parasites generated using gRNA 1 and gRNA2 were both positive for pCSC1-HAglmS integration (Figure S8B). To confirm expression of HA tagged *Pf*CSC1, a Western blot was performed, but no signal could be detected with HA antibodies. Immunofluorescence microscopy of CR-CSC1(WT) parasites detected very weak labelling with HA antibodies at the parasite periphery in the trophozoite stage (Figure S8C). Given the lack of co-localisation with EXP2 at schizont stages, this HA labelling is most likely at the parasite membrane rather than PV. In schizonts, HA-tagged CSC1 appeared to reside within internal structures possibly membranes of the newly forming merozoites while EXP2 remained predominantly at the PVM around the schizont’s periphery (Figure S8C).

To determine if *Pf*CSC1 was important for parasite growth, similar to *Pf*ROM8, we attempted to knock it down by following addition of 2.5 mM GlcN. As we could not detect the protein by western blot qRT-PCR was then performed on CR-CSC1(WT) parasites. The *csc1* mRNA levels of parasites were grown in the presence of 2.5 mM GlcN were ∼30% relative to no GlcN (∼70% knockdown of *csc1* mRNA) whilst as expected, levels of *exp2* mRNA remained unchanged in 2.5 mM GlcN (Table S5).

We next assessed the impact that CSC1 knockdown had on parasite growth. For this, early ring stage (0-4 h) stage parasites in cycle 0 (C0) were treated with 2.5 mM GlcN or left untreated. Measurement of parasite survival after 8 days in culture was assessed using a SYBR Green assay, which revealed knockdown with GlcN was significantly detrimental to parasite growth (Figure 5A). Parasite growth was also assessed by visualization of Giemsa-stained smears and compared to untreated parasites, where a delay in parasite growth was evident by trophozoite stage in the cycle following knockdown (C1)(Figure S9A-C). The delayed growth phenotype was not observed in all parasites, however, likely because knockdown was only partial. As a result, a proportion of the populations were able to survive, replicate and re-invade new RBC.

**Figure 5:**
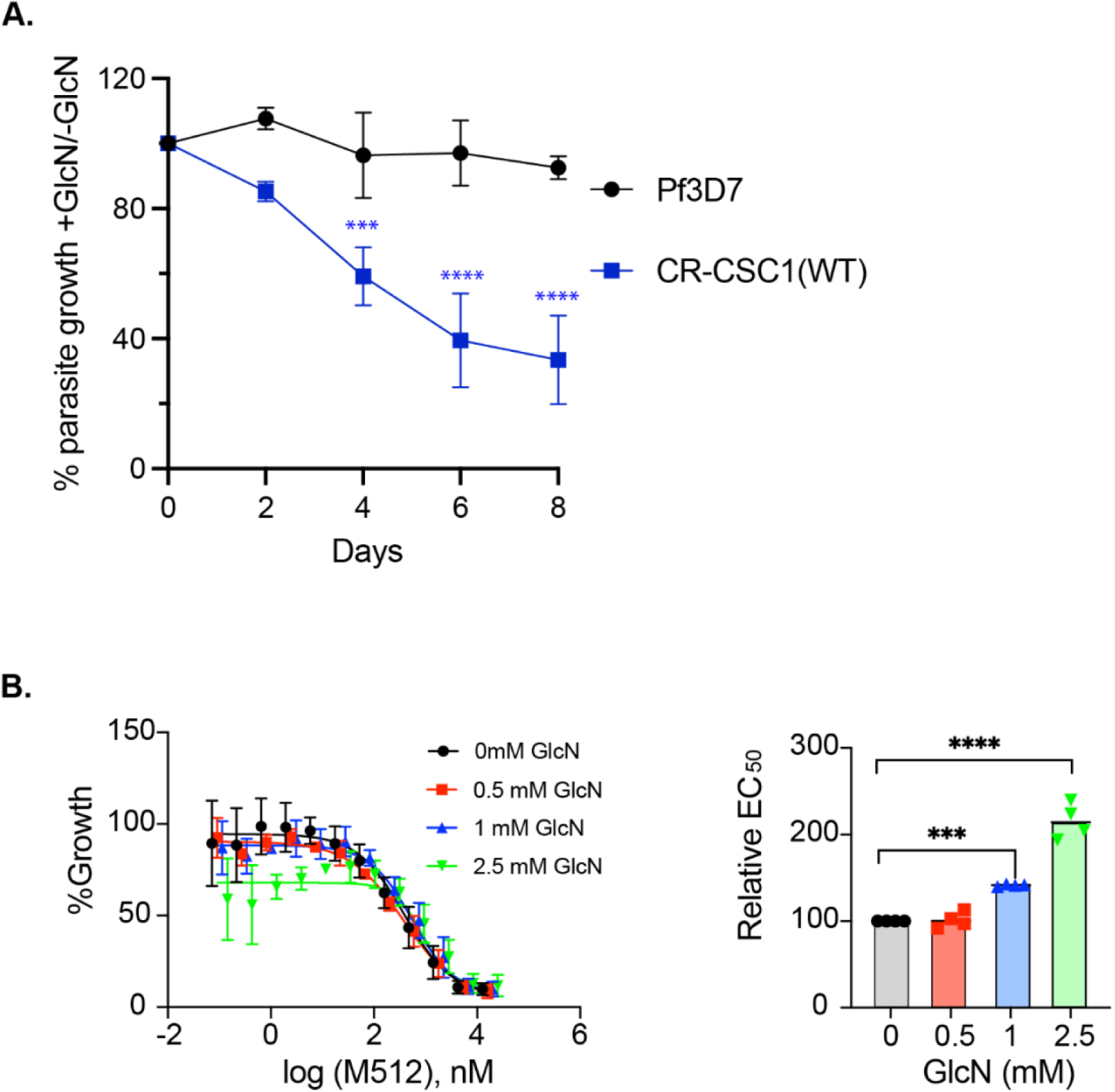
Depletion of *Pf*CSC expression leads to perturbed parasite growth *in vitro*. (**A**). Parasite survival of *Pf*CSC-HAglmS after treatment with GlcN compared to untreated parasites as determined by SYBR Green assay. Shown is the mean ± standard deviation (n= 3). Statistical significance was analysed using 2-way ANOVA comparing parasite lines at each timepoint, followed by Sidak’s multiple comparisons test (*** *p* ≤ 0.001; \*\*\*\**p* ≤ 0.0001). **B)** Effect of M-512 on CR-CSC(WT) parasites grown in the presence or absence of different concentrations of GlcN over 72 h as determined by SYBR Green assay (left graph) and the change to EC_50_ relative to parasites grown without GlcN (Right graph, n=4 biological repeats and a one-way ANOVA with Dunnett’s multiple comparison test was used. **p<0.001 and ****p<0.0001..

Using the CR-CSC1(WT) parasites, we next examined the impact of CSC1 knockdown on the EC_50_ of M-512. Ring-stage parasites were cultured in 0, 1 or 2.5 mM GlcN and the following cycle, M-512 was added. A standard 72-hour growth inhibition assay was then undertaken to determine the EC_50_. (Figure 5, left). Surprisingly, knockdown of CSC1 led to more than a 2-fold increase in the EC_50_ (Figure 5, right).

## Discussion

M-512 was identified in a screen of the MMV Pathogen Box as an inhibitor of parasite invasion. Whole genome sequencing of three M-512 resistant populations recovered two different mutations in *pfrom8* and a double mutation in *pfcsc1* and *val trna lig*. To corroborate the *Pf*ROM8 mutations were responsible for resistance to M-512 we introduced these mutations into wildtype parasites but could only recover the *Pf*ROM8(L562R) mutant. The *Pf*ROM8(L562R) mutation appeared slightly deleterious for growth, suggesting the mutation reduced normal functioning of the protease. When *pfrom8* was modified, the gene was also tagged with *glmS* ribozyme and knockdown of *Pf*ROM8 appeared to reduce parasite growth particularly in the L562R mutant. Following *Pf*ROM8 knockdown, intermediate concentrations of M-512, could partly restore parasite growth indicating the compound was acting as a *Pf*ROM8 agonist.

We did not attempt to introduce the *Pf*CSC1(L954F) mutation into wildtype parasites as we were unsure whether it might also need the Val tRNA Ligase(M1061I) mutation. We were able to append a *glmS* tag and to knock down expression, which like CR-ROM8(WT) led to reduced parasite growth and increased resistance to M-512. We are confident, that *Pf*CSC1 is involved in resistance to M-512 as related work has shown that resistance selection to a more potent analogue of M-512 (WEHI-2047326) also selects for different mutations in *Pf*CSC1 and *Pf*ROM8 (Nguyen, Boulet et al. 2025).

M-512 was identified as an inhibitor of invasion during our egress and invasion assay which spans a four-hour period (Dans, Weiss et al. 2020). During this time merozoites complete their development within the schizont, egress, invade new erythrocytes and then transform into intracellular ring-stage parasites. To determine if M-512 was a specific invasion inhibitor, purified merozoites were exposed to the compound for 30 minutes during invasion. In this plate-based assay M-512 exhibited modest inhibitory activity but was not as potent as other inhibitors that target actin1/profilin and *Pf*START1 (Dans, Weiss et al. 2020, Dans, Piirainen et al. 2023, Dans, Boulet et al. 2024). Since other members of the rhomboid protease family namely ROM1 and ROM4, are implicated in helping to shed the merozoite’s surface coat during invasion we initially believed *Pf*ROM8 could have a similar invasion role. Since *Pf*ROM8 is predicted to be an active protease as it possesses most of the functionally important motifs typical of these proteases (Ha, Akiyama et al. 2013), we performed live cell invasion imaging in the presence of M-512. Here we could see no significant effects on invasion, indicating that M-512 may be exerting its influence on the development of the merozoites, which was recently shown to stall when *rom8* was disrupted using the inducible frame-shift SHIFTiKO method (Ramaprasad and Blackman 2024). M-512 was also observed to be a mid-level inhibitor of ring and trophozoite stage parasites, suggesting the compound may exert a broad effect on intraerythrocytic forms by targeting *Pf*ROM8 and *Pf*CSC1 (Dans, Weiss et al. 2020).

*Pf*CSC1 belongs to a family of osmosensitive or mechanosensitive calcium-permeable cation channels conserved across eukaryotes (Hou, Tian et al. 2014). These channels help cells respond to certain biotic and abiotic stresses and in the plant *Arabidopsis*, the At4G22120 channel responds to hyperosmotic shock by permitting conductance of Ca^2+^, K^+^, and Na^+^ ions across the plasma membrane, leading to the channel being named Calcium permeable Stress-gated cation Channel 1 or *At*CSC1. In *Arabidopsis*, a related group of mechanosensitive *At*OSCA channels have been shown to form homodimers (Zhang, Shan et al. 2023). From this structure we modelled the *Pf*CSC1 dimer, which showed that the *Pf*CSC1 L954F mutation is predicted to be located near the parasitophorous vacuole-exposed region of the membrane protein (Figure 1C). As the *Pf*ROM8 mutations are likewise predicted to reside near the PV-exposed region of the protease, it is proposed that *Pf*ROM8 may interact with *Pf*CSC1, with the resistance mutations lining a PV-exposed pocket or region where M-512 may bind. The compound could therefore act as an agonist by mimicking the mechanosensitive or osmosensitive deformations exerted on the *Pf*CSC1 channel and lock it in an open conformation. Mutations lining the M-512 binding pocket may reduce the pocket’s affinity for M-512, conferring resistance to the compound’s effects. At this stage we do not know if *Pf*ROM8’s proteolytic activity is required but there are examples of rhomboid proteases regulating the activity of Ca^2+^ ion channels of a different family in human cells (Grieve, Yeh et al. 2021).

Our companion study described improving the potency of M-512 through medicinal chemistry to produce analogue WEHI-2047326 (WEHI-326), which had single digit nanomolar potency (Figure S1A) and very little toxicity to human HepG2 cells (>40 CC_50_) (Nguyen, Boulet et al. 2025). WEHI-326 remained on target as our M-512 resistant parasites were also resistant to the potent analogue (Nguyen, Boulet et al. 2025). Resistance selection to WEHI-326, followed by whole genome sequencing, also revealed mutations in *Pf*ROM8 and *Pf*CSC1, indicating that WEHI-326 shares the same mechanism of action as M-512. WEHI-326 was also subjected to a AReBAR resistome pool study (Bower-Lepts, Rawat et al. 2024) where pools of barcoded parasites, each containing a known resistance mutation, were cultured with WEHI-326. Parasites containing a L800P mutation in *Pf*CSC1, originally selected with MMV Pathogen Box compound MMV407834 (Figure S1A), were the only parasites that could proliferate in WEHI-326, again supporting *Pf*CSC1 as being a major target (Nguyen, Boulet et al. 2025). Adding to the appeal of this series as a seasonal transmission blocking treatment was that WEHI-326 maintained potency towards stage II-III and stage V gametocytes as well as female and male gametes (Nguyen, Boulet et al. 2025). It also strongly blocked the formation of oocysts in mosquito guts following membrane feeding assays (Nguyen, Boulet et al. 2025).

WEHI-326 was, however, susceptible to metabolic breakdown in liver microsomes and this probably resulted in the failure of the compound to reduce the parasitemia of mice infected with *Plasmodium berghei* (Nguyen, Boulet et al. 2025). WEHI-326 also has an minimum inoculum of resistance (MIR) of 4 x 10^7^, which indicates a moderate risk of parasite resistance arising to the compound (Nguyen, Boulet et al. 2025). This agrees with our own experiments where parasite resistance to M-512 was observed in all five compound-selected populations. The rate of kill of WEHI-326, also known as parasite reduction ratio, was similar to pyrimethamine and therefore slower than the rapidly acting artemisinin compound (Nguyen, Boulet et al. 2025).

## Conclusions

We have demonstrated that M-512’s mechanism of action is likely to agonistically activate *Pf*ROM8 and *Pf*CSC1, allowing the unrestricted flow of cations across the parasite’s plasma membrane and dysregulation of ionic homeostasis, which in turn inhibits parasite viability. A major future goal is to discover if these two proteins interact and are functionally cooperative, and if this mechanism involves proteolytic activity. It will also be interesting to unravel what the natural function of the putative *Pf*ROM8/*Pf*CSC1 complex is and why it is important for parasite growth. Other future challenges will be to engineer metabolically stable analogues M-512 that retain their potency and are less susceptible to parasite resistance.

## Materials and Methods

### P. falciparum culture

*Plasmodium falciparum* 3D7 and W2mef parasites and transfectants were cultured in human red blood cells provided by the Australian Red Cross Lifeblood, at 4% haematocrit and maintained at 37°C in a special gas mixture (1% O_2_, 5% CO_2_, 94% N_2_)(Trager and Jensen 1976). The medium used was complete RPMI medium: RPMI-1640 (Sigma), 25 mM HEPES (GIBCO), 0.37 mM hypoxanthine (Sigma), 31.25 μg/mL gentamicin (GIBCO), 0.2% NaHCO_3_ (Thermo Scientific), 0.5% AlbuMAX II (GIBCO).

### Live cell imaging

Synchronous W2mef parasites at the late schizont stage were placed into each well (0.2 mL of 0.08% haematocrit) of an eight chambered coverslip (Ibdi) and mounted in a humidified environmental chamber at 37°C and gassed (5% C0_2_, 1% O_2_ and 95% N_2_). This was observed in a Zeiss Axio Observer microscope with a LCI Plan-Neofluar 63x/1.3 Imm Corr DIC M27 objective.

### Resistance selection to M-512

5 mL of culture containing 1x10^8^ parasites in 4% haematocrit (HCT) was added to each well of a 6-well plate. 5 wells labelled A - E were treated with 5 µm MMV020512 (M-512) until the parasites began to die, as observed by Giemsa-stained thin blood smears. A 6^th^ control well of parasites was treated with 0.1% DMSO and was maintained to monitor parasite health. M-512 was then removed to allow the parasite population to recover, and the process was repeated a total of three times. 72-hour growth inhibition assays were then performed (see below) to determine if the populations had become resistant. If so, clonal lines were derived by diluting the parasites into 96-well plates at 0.3 parasites/well.

### Growth inhibition assays

M-512 was serially diluted and added to ring-stage parasites at 0.3% parasitaemia in 2% HCT with an uninfected RBC control. The parasites were grown for 72-hours in complete RPMI as described above and were then frozen. When applicable, 0.25 mM or 2.5 mM GlcN was also added. Parasite lactate dehydrogenase activity was measured using a modified protocol from (Makler and Hinrichs 1993). 30 µL of thawed parasites were added to 75 µL of LDH reagent (0.1 M Tris pH 9.0, 20 g/L lactic acid, pH 7.5, 0.2% Triton-X100, 0.5 g/L acetylpyridine adenine dinucleotide (APAD, Sigma), 200 μg/mL nitroblue tetrazolium (NBT; Sigma), 1 μg/mL phenozine ethosulfate (PES; Sigma)). After monitoring for a LDH-dependent purple colour change, absorbance at OD650 was measured and growth inhibition curves were plotted in PRISM V10 as per (Dans, Boulet et al. 2024).

### Whole genome sequencing

After clonal lines had been established from the M-512 resistant parasite populations, their genomic DNA (gDNA) was extracted (DNeasy Blood and Tissue kit (Qiagen)). DNA was prepared for sequencing first by constructing a library using the native barcode sequencing kit (SQK-LSK109) as per manufacturer’s instructions (Oxford Nanopore Technologies, UK). Two sequencing runs were performed splitting up the samples, 4 in one sequencing run and 3 in the other to optimise data generation. These runs were performed on a MinION platform with MIN111/R10 Flow cells and MinIT (Software 18.09.1) to generate fastq files. Post sequencing, fastq files were base called and demultiplexed using (Guppy V2.3.1). Reads were aligned against the *P. falciparum* 3D7 reference genome (PlasmoDB version 46) using minimap2 (V2.21) with default parameters. Alignment files were processed with samtools utilities (V1.9) to generate sorted bam files. Genotype likelihoods were then computed using bcftools mpileup (V1.9), using the skip indels parameter and only retaining bases with quality 7 or above. Variant calling was conducted using bcftools multiallelic-caller (V1.9) to generate haploid SNP candidates for each sequenced isolate. Candidate SNPs in blacklisted genomic regions (subtelomeric, centromeric, or hypervariable regions) were removed using bedtools subtract (V2.29.2) to only retain SNPs in the core genome. Quality filtration of SNP calls, with a minimum depth of coverage of 10X and at least 70% of reads to support the called allele, was performed using custom awk script. Filtered candidate SNPs for each isolate were then imported into R statistical software (V3.6.1) and overlaps between the 3D7 WT and resistant isolates were explored. SNPs present in the 3D7 WT isolate were removed, accounting for differences between 3D7 WT isolate and cultured 3D7 isolate. Functional annotation of non-WT candidate SNPs was performed using the VariantAnnotation package (V1.32.0), retaining only nonsynonymous SNPs. The annotation information including biological annotations, including gene ontology terms and expression profiles, if available were collated to identify causal resistance candidate SNPs using an in-house R package (PlasmoCavalier). Genomic sequencing data is available from the European Nucleotide Archive; accession number (submission pending).

### PCR validation of resistance mutations

To ensure the SNPs detected in *pfrom8*, *pfcsc1* and *val-trna lig* were genuine, PCRs were performed using gene specific Mut_F and Mut_R primer sets (Table S2), gDNA from the clonal lines and Phusion DNA polymerase as per manufacturer’s instructions (Thermo). The PCR products were inserted into plasmid using the CloneJET PCR cloning system (Thermo) and sequenced with plasmid specific primers. Sanger sequencing was performed (Micromon, Monash University) and the data assembled using SnapGene V5.3.3.

### Donor plasmid construction and transfection for ROM8

A 947 bp region within the 5’ half of ROM8 was amplified using ROM8_1F and ROM8_2R (Table S2) from 3D7 gDNA using Phusion DNA polymerase as per manufacturer’s instructions (Thermo). A 648 bp region of corresponding to the 3’ half of *rom8* was synthesised as gBlocks using a *Saccharomyces cerevisiae* codon bias (IDT). The annealed gBlocks were insert into plasmid using the CloneJET PCR cloning system (Thermo) which served as a PCR template using primers ROM8_3F and ROM8_4R (Table S2). The native and recodonised regions of ROM8 were joined by overlapping PCR to form the ROM8 homology block that was inserted into the p1.2 plasmid via *BglII* and *SpeI* sites (Marapana, Dagley et al. 2018, Dans, Piirainen et al. 2023). A 514 bp fragment of the 3’ half of *rom8* was amplified by ROM8_EcoR1.2 and ROM8_6R (Table S2) and inserted into the *EcoRI* and *KasI* sites of plasmid p1.2 to form the ROM8 3’ homology block. In this CR-ROM8(WT) donor plasmid, the 5’ and 3’ *Pf*ROM8 homology blocks were separated by a human dihydrofolate reductase gene cassette to select for repair and replacement of *rom8* by double recombination using compound WR99210 (Jacobus Pharmaceutical Company) (Figure S4A). Recombination and repair with the donor plasmid was facilitated by excision of the native *rom8* locus with recombinant Cas9 complexed with synthetic *rom8* specific guide RNA (Table S2) (McHugh, Bulloch et al. 2023).

### Donor plasmid construct and transfection for CSC1

To create the targeting construct pCSC-HAglmS for conditional knockdown of the *P. falciparum csc1* gene (PF3D7_1250200), a 5’*csc1* homology block was cloned into the *BglII* and *PstI* sites of pRhopH2_HAglmS plasmid and a 3’ *csc1* homology block was cloned into the *StuI* and *EcoRI* sites. The 5’homology block was synthesised by Invitrogen GeneArt services; it comprised of 1094 base pair (bp) of sequence upstream of the *csc1* stop codon, of which the last 426 bp was recodonised and excluded the final intron. This resulted in fusion of the CSC1 C-terminus with a triple hemagglutinin (HA) and single streptavidin epitope tag. The 3’homology block was PCR amplified from *P. falciparum* genomic DNA using DO1500/DO1501. A total of 100 µg of the pCSC-HAglmS construct was transfected together with a Cas9 ribonucleoprotein complex, as described previously for ROM8.

### Growth rate assays

Synchronised parasites were dispensed in duplicates, in two 96-well plates, at parasitemia ∼0.3-0.5% trophozoite, 4% hematocrit, in the presence of 1 µM M-512 or 0.01% DMSO when applicable. The first plate’s OD650 was analysed (LDH readout) immediately (corresponding to time 0 h of cycle 1), while the second plate was incubated for 48 h in normal culturing conditions. After the 48 h, LDH activity of the culture was read at OD650 (time 48 h of cycle 1), while the remainder was diluted twice (between 1:4 to 1:10 dilution) with LDH activity of the first dilution measured right away (time 0 h of cycle 2), and the second dilution was measured 48 h later (time 48 h of cycle 2). The OD650 of technical duplicates was averaged for each timepoint and each condition, and for each cycle the ratio OD650(48 h)/OD650(0 h) was calculated.

### Conditional knockdown of CR-ROM8 and PfCSC1 expression in P. falciparum and growth analysis

Heparin synchronized CR-CSC1(WT) parasites were treated in cycle 0 (C0) with 2.5 mM GlcN at 0-4 h post-invasion; untreated parasites and 3D7 parasites treated with 2.5 mM GlcN served as controls. Parasite growth was visualised for the next two cycles following GlcN treatment (C1, C2) via Giemsa-stained blood smears, with images taken with a SC50 5-megapixel or IX71 color camera (Olympus). Each cycle, 200 μL aliquots of resuspended RBC were taken and stored at –20°C for SYBR Green I assay. Briefly, 30 μL of freeze-thawed suspension was added in triplicate to 96-well assay plates (Costar) to which an equal volume of Lysis buffer (20 mM Tris pH 7.5, 5 mM EDTA, 0.008% saponin (w/v) & 0.008% Triton x-100 (v/v)) (Smilkstein, Sriwilaijaroen et al. 2004) containing 0.2 µL/mL SYBR Green I Nucleic Acid Gel Stain (10,000x in DMSO; ThermoFisher) was added to each well. Following a 1 h incubation in the dark at RT, fluorescence intensity was read on a Glomax® Explorer Fully Loaded (Promega) with emission wavelengths of 500-550 nm and an excitation wavelength of 475 nm. Graphs were generated using GraphPad Prism (V.8.4.1). Three biological replicate experiments were performed, and a two-tailed unpaired t-test was used to determine statistical significance between groups. Data is presented as the mean and error bars representative of standard deviation.

### Western blot analysis of ROM8 parasites

CRISPR ROM8 parasites (WT, L562R and T662K) schizonts were lysed 10 min on ice with 0.1% saponin in PBS with Protease Inhibitors cocktail (Roche; PBS+PI). Parasites were washed with PBS+PI, resuspended in 10 x volume of non-reducing sample buffer (NRSB; 50 mM Tris-HCl pH 6.8, 2 mM EDTA, 2% SDS, 10% glycerol, 0.005% phenol blue), sonicated 3 × 30 seconds (Diagenode sonicator) and boiled 10 min at 80 °C. Samples were run on a pre-cast 4-12% NuPAGE Bis-Tris gel (Invitrogen), transferred on a nitrocellulose membrane using iBlot (Invitrogen). Blocking and antibodies were in 3% BSA in TBST. Primary antibodies anti-HA (mouse, Sigma; 1:1000) and anti-START1 (rabbit, WEHI (1:1000)(Dans, Boulet et al. 2024) were incubated overnight at 4°C; secondary antibodies (Invitrogen rabbit-Alexa Fluor Plus 680 and mouse-Alexa Fluor Plus 800; 1:10,000) 1 h at room temperature. Fluorescence was measured using an Odyssey imaging system.

### Immunofluorescence Microscopy

Thin blood smears of asynchronous CR-ROM8(WT / L562R / T622K) were fixed 5 min in ice-cold 90% acetone / 10% methanol. Samples were blocked and antibodies prepared in 3% BSA, 0.02% TX100 in PBS. Primary antibodies used were anti-HA (mouse, Sigma), combined with either anti-MSP1-19 (rabbit), anti-EXP2 (rabbit), or anti-RhopH3 (rabbit), all 1:500 dilution (Table S4). After an overnight incubation at 4°C, samples were thoroughly washed in 0.02% TX100 in PBS, and secondary antibodies added for 1 h at 1:2000 dilution, with 5% goat serum: anti-mouse-Alexa Fluor 594 (Invitrogen) and anti-rabbit Alexa Fluor 488 (Invitrogen). Following washes, VectaShield Antifade Mounting Medium (with DAPI, H-1200) was added, coverslip placed and sealed, before imaging on a Zeiss Cell Axio Observer (Carl Zeiss). Images were processed with ImageJ.

### Quantitative reverse-transcription polymerase chain reaction (qRT-PCR)

Gene expression was examined in CR-ROM8(WT), CR-ROM8(L562R) and CR-CSC1(WT) parasites after being grown in the absence or presence of 2.5 mM GlcN for 1 cycle for CR-ROM8 parasites and 2 cycles for CSC1(WT). Briefly, RBCs infected with trophozoite-stage parasites were harvested by lysing with 0.15% saponin in PBS. Following RNA extraction using the RNeasy Mini Kit (Qiagen), cDNA was synthesised using QuantiTec cDNA synthesis kit (Qiagen) and then subjected to qRT-PCR using 2×SensiFASTMix™ SYBR® Low-ROX master mix (Bioline). The following primer sets were used: DO2079/DO2081 for CR-*rom8*, DO1749/DO1772 for *csc1*, DO1812/DO1813 for *exp2* (PF3D7_1471100), DO1810/DO1811 for the house-keeping gene fructose-bisphosphate aldolase (fbpa;PF3D7_1444800) (sequences provided in Table S3). The expression level of *csc1* and *exp2* were normalised against the *fbpa* house keeping gene and the fold-change calculated using the 2ΔΔCt method (Rao, Huang et al. 2013).

## Acknowledgments

We thank Lifeblood Biological Resources Australia for providing the human red blood cells. This work was supported by the Victorian Operational Infrastructure Support Program received by the Walter and Eliza Hall and Burnet Institutes. This work was funded by the National Health and Medical Research Council of Australia (Ideas Grant to W.N. and P.G. 2001073 and Ideas Grant to J.M, N.C and T.D.K-W 1182000). B.E.S. is a Corin Centenary Fellow.

