## Supplementary data for "Aryl N-acetamide compounds exert antimalarial activity by acting as agonists of rhomboid protease *Pf*ROM8 and cation channel *Pf*CSC1"

**Title.**

**Table S1. Whole Genome Sequencing variants from M512 resistant genomes.**
This is a separate Excel file.

### Table S2. DNA primers, sequences and guide RNAs used for ROM8 in this study.

| **Primer name** | **Sequence** | **Purpose** |
|---|---|---|
| ROM8_Mut.F | ATGATTCCTTAAAAAATTTTGGTAGTAATTTTCCACA | Amplifies a region of *rom8* that encompasses the L562R and T622K mutations. |
| ROM8_Mut.R | TGCACACTAATACTAATTTTGTCTACATTAAAATTAG<br>TCATCA | |
| CSC_Mut.1F | GCTTTTTTCAGTGGCCTTCTTTCAAGTA | Amplifies a region of *csc1* that encompasses the L954F mutation. |
| CSC_Mut.2R | TGTTTCGGCCGACTGATATGAACT | |
| Val.tLig_Mut.F | GTACAATCAAGTGAAGAATACCTTAAACCATTATTAA<br>GTAGT | Amplifies a region of *Val tRNA Ligase* that encompasses the M1061I mutation. |
| Val.tLig_Mut.R | CTATGCATTCTTTAAACACTCATTTATTTCACATATTA<br>TATTCGA | |
| ROM8_IntF | GACGGTGTGGGGGATTATGGTGATGATGA | Primes outside integration block to check if *rom8* locus has been modified. |
| Rom8_1F | AGATCTACAGCTGTAGTAGTTTTACGAGGAGAGGA | Used with Rom8_2R to amplify the 5' native homology block. |
| Rom8_2R | TATAACCCACTGAATAAATGTCACATATAACGTTAAA<br>CT | |
| Rom8_3F | TGACATTTATTCAGTGGGTTATATTTATCCTGTTAAT<br>TAGCGT | Used with Rom8_4R to amplify the recodonised ROM8 homology block. |
| Rom8_4R | ACTAGTTTTAGTAGTCCGGGACGTCGTACGGGTAA<br>GAGGCTGCAGCATTGGGACAATTTTGGACAACATA<br>GGT | |
| HAglmS_R | AGATCATGTGATTTCTCTTTGTTCA | Primes from glmS tag in p1.2 donor plasmid with upstream primer ROM8_IntF to check for correct integration into *rom8*. |
| ROM8_EcoR1.2 | GAATTCACCGAAAGCAGTAGTAGTTCTTCAGGAATA<br>ATTGGA | This primer pair amplifies the 3' homology block for insertion into the p1.2_Rom8 donor plasmid by *EcoR1* and *KasI* sites. |
| Rom8_6R | GGCGCCTTAATTTGGACAATTCTGAACAACATAGGT<br>GGA | |
| Recodonised 3' flank of *rom8*:<br>• L562R mutant (orange, in brackets)<br>• T622K mutant (purple, in brackets) | TTTATCCTGTTAATTAGCGTTAAATCTGACTTTCCCCTGACTCCGAGCAACGATTCACTTA<br>AAAATTTCGGGTCAAACTTCCCGCATAATATCTTcAAAAATGCAGAGTTCTATAGG***TTG(C***<br>***GC)***TTTACTTCTCTTTTCCTACACTCCAACTTTAACCACATTTGCGCGAATACTTATGTTCA<br>GCTGACAGTGGGCTTCCTGTTAGAGTACCTGTATGGCACTTATGTAGTGTTTTTGGTTTA<br>CGTTTTTACGGGGATCTATGGTATAATTTTGAGCAGTCCACTTACTTACTGTTATAGT***ACG***<br>***(AAG)***ACTGAGTCCTCTTCTTCTAGTTCTGGGATTATTGGGATCTTTTTCTCCGAGATTCTT<br>ATGATGACCAATTTCAACGTAGATAAAATCTCTATAAGTGTCCATCTTTTTTGCTTCTTCC<br>TGTTACTGCTTTTTTTGAAGTTCTCTTTAAACACTGTTAGTATCAATATTTATAGCCATTTT<br>TTCGGATTTCTGGGCGGATTTCTAATAGGAATCATCTTAAAGAGGAACCAGTTGAAGTAT<br>TTCTTAAGAAATAATTTTTTGATACAAATACTTTGTTTCGTCTTTCTGGTAATCAGTCTAGG<br>CGCAGCCATCTTCATTAGCACCTATGTTGTCCAAAATTGTCCCAAT | |
| gRNA | TCTTTCGAGTGAAAAGCAGA<br><br>rUrC rUrUrU rCrGrA rGrUrG rArArA rArGrC<br>rArGrA rGrUrU rUrUrA rGrArG rCrUrA rUrGrC rU | Guides the recombinant Cas9 enzyme to cut the native *pfrom8* locus (but not the recodonised *pfrom8* DN<br>A template) |
| DO2079 qPCR F | TCGAATCTTCCCGCAATTCTC | Primers used to amplify *pfrom8* by qRT-PCR. |

| | | |
|---|---|---|
| DO2081 qPCR R | CGCGCAAATGTGGTTAAAGT | |
| DO1810_Fructose-BPA-F | TGTACCACCAGCCTTACCAG | Primers used to amplify PfFructose-BPA (Pf3D7_1444800) by qRT-PCR |
| DO1811_Fructose-BPA-R | TTCCTTGCCATGTGTTCAAT | |

**Table S3. DNA primers, sequences and guide RNAs used for CSC1 in this study.**

| Primers | Sequence | Purpose |
|---|---|---|
| DO276 | GTGATTTCTCTTTGTTCAAGGA | GlmS R to check 5’ integration |
| DO1494 | aaaaagtatagggaccctag | CSC gRNA1 |
| | rArA rArArA rGrUrA rUrArG rGrGrA rCrCrC rUrArG rGrUrU rUrUrA rGrArG rCrUrA rUrGrC rU | |
| DO1496 | cgaacaaatttggatagtca | CSC gRNA2 |
| | rCrG rArArC rArArA rUrUrU rGrGrA rUrArG rUrCrA rGrUrU rUrUrA rGrArG rCrUrA rUrGrC rU | |
| DO1498 | GGTCATGTATTCTGATGAAATCGG | CSC Homology Box (HB) 1 F |
| DO1500 | CGAgaattcaggcctCATATATATGTGTGTACATATGTG | CSC HB 2F |
| DO1501 | CGAgaattcaggcctCATATATATGTGTGTACATATGTG | CSC HB 2R |
| DO1668 | GTGAACACACGTACGTGTTAC | CSC integration 5’ end |
| DO1749 | GAAATCAGTGCGTGGACTTTTG | CSC F for qRT-PCR |
| DO1772 | CCATGTGAGTCTAAGTTGGTAC | CSC R for qRT-PCR |
| DO1810 | TGTACCACCAGCCTTACCAG | FBPA F for qRT-PCR |
| DO1811 | TTCCTTGCCATGTGTTCAAT | FBPA R for qRT-PCR |
| DO1812 | GTGGTGGGTATTGTTAGTAAGAG | EXP2 F for qRT-PCR |
| DO1813 | GTGGCAAAGTTGTTTCTGCATTC | EXP2 R for qRT-PCR |

**Table S4. Antibodies used in this study**

| **Antibody** | **Raised in:** | **Provider** | **Dilution** (WB: western blot; IFA: immunofluorescent assay) |
|---|---|---|---|
| HA | Mouse | Sigma | 1:1000 (WB)<br><br>1:500 (IFA) |
| *Pf*START | Rabbit | WEHI (see Methods) | 1:1000 (WB) |
| *Pf*EXP2 | Rabbit | WEHI (PMID: 19536257) | 1:2000 (WB)<br><br>1:500 (IFA) |
| *Pf*MSP1-19 | Rabbit | WEHI (PMID: 18333885) | 1:2000 (WB)<br><br>1:500 (IFA) |
| *Pf*RhopH3 | rabbit | WEHI (PMID: 28252383 | 1:300 (IFA) |
| Rabbit-Alexa Fluor Plus 680 | Goat | Invitrogen | 1:10000 (WB) |
| Mouse-Alexa Fluor Plus 800 | Goat | Invitrogen | 1:10000 (WB) |
| Rabbit-Alexa Fluor 488 | Goat | Invitrogen | 1:2000 (IFA) |
| Mouse-Alexa Fluor 594 | Goat | Invitrogen | 1:2000 (IFA) |

**Table S5. Expression of *pfcsc1* and *pfexp2* relative to treatment with glucosamine**

| Parasite line | PfCSC1 | PfEXP2 |
|---|---|---|
| CR-CSC1(WT)-G1 | 28.4% | 101.3% |
| CR-CSC1(WT)-G2 | 31.2% | 99.2% |

**A.**

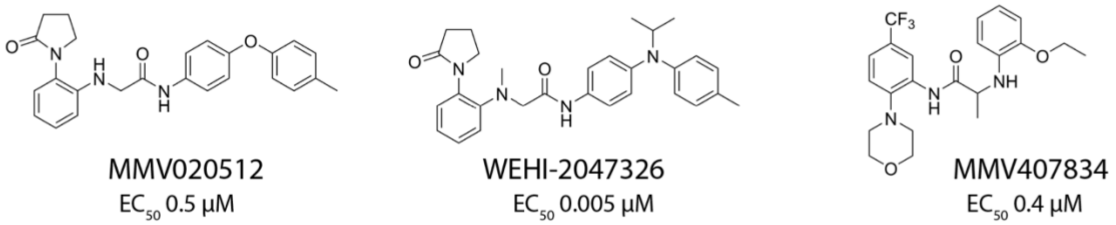


**B.**

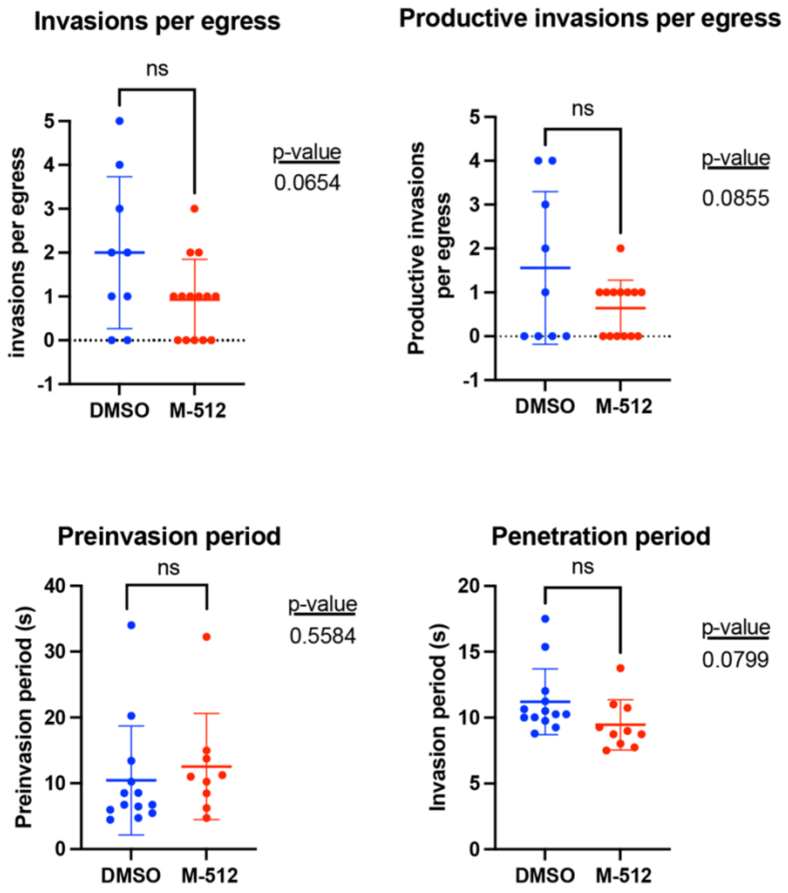


**Figure S1. M-512 did not significantly reduce the invasion efficiency of *Plasmodium falciparum* merozoites into human erythrocytes when observed by live cell imaging. A)** Structures of inhibitory compounds discussed in this study. The $EC_{50}$ values of 3D7 parasites challenged with these compounds as presented by Medicines for Malaria Venture. **B).** Late stage schizonts were treated with 5µM M-512 or 0.05% DMSO before being observed until the merozoite egressed and invaded nearby erythrocytes. The number of invasions per egress, the number of egressed merozoites that became ring-stage parasites (productive invasions), the time from first merozoite on erythrocyte contact until penetration (preinvasion) and the penetration times were measured. Means and SD are indicated, and p-values were calculated using unpaired t-tests in PRISM V10.

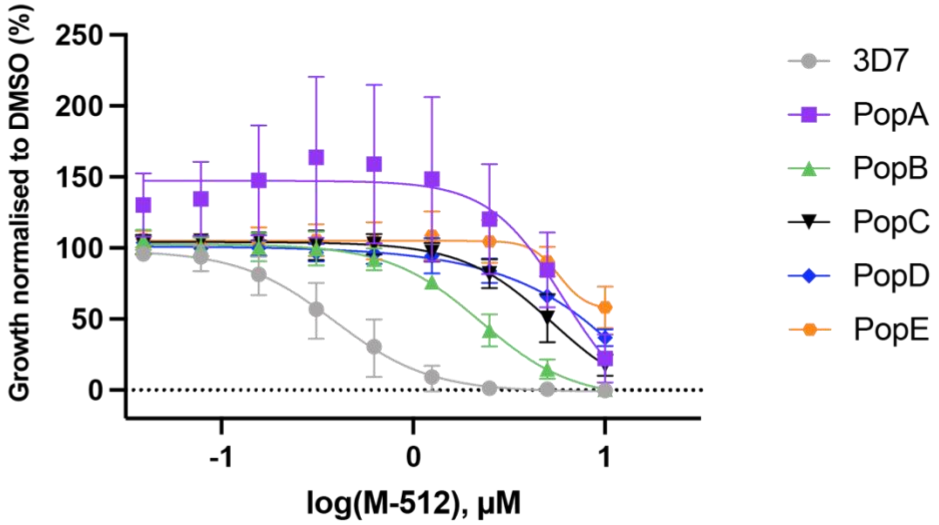


| | 3D7 | PopA | PopB | PopC | PopD | PopE |
|---|---|---|---|---|---|---|
| **EC50** | 0.388 | 6.051 | 2.243 | 5.301 | 39.390 | 5.745 |
| **95% CI (Top)** | 87.56 to 115.4 | 128.4 to 171.4 | 97.74 to 108.0 | 99.51 to 109.4 | 96.69 to 106.4 | 100.0 to 110.5 |

**Figure S2. Parasites selected for resistance to MMV020512 have mutations in the genes for Rhomboid protease 8 (PF3D7_1411200) and Calcium Stress Response (CSC1, PF3D7_1250200).** Growth of five populations independently selected for resistance to MMV020512 (M-512). Parasites were challenged with M-512 for 72-hours followed by measurement of lactate dehydrogenase (LDH) as a proxy for growth. The data points are means of three biological replicates, each of two technical replicates normalised to their respective DMSO vehicle controls. The $EC_{50}$ for PopE is underestimated as the inhibition curve does not fall below 50% at the highest drug concentration (10 µM). The whiskers indicate SD. Below the graph are $EC_{50}$ values and 95% top CI range values. Graphs and analysis were performed in PRISM V10.

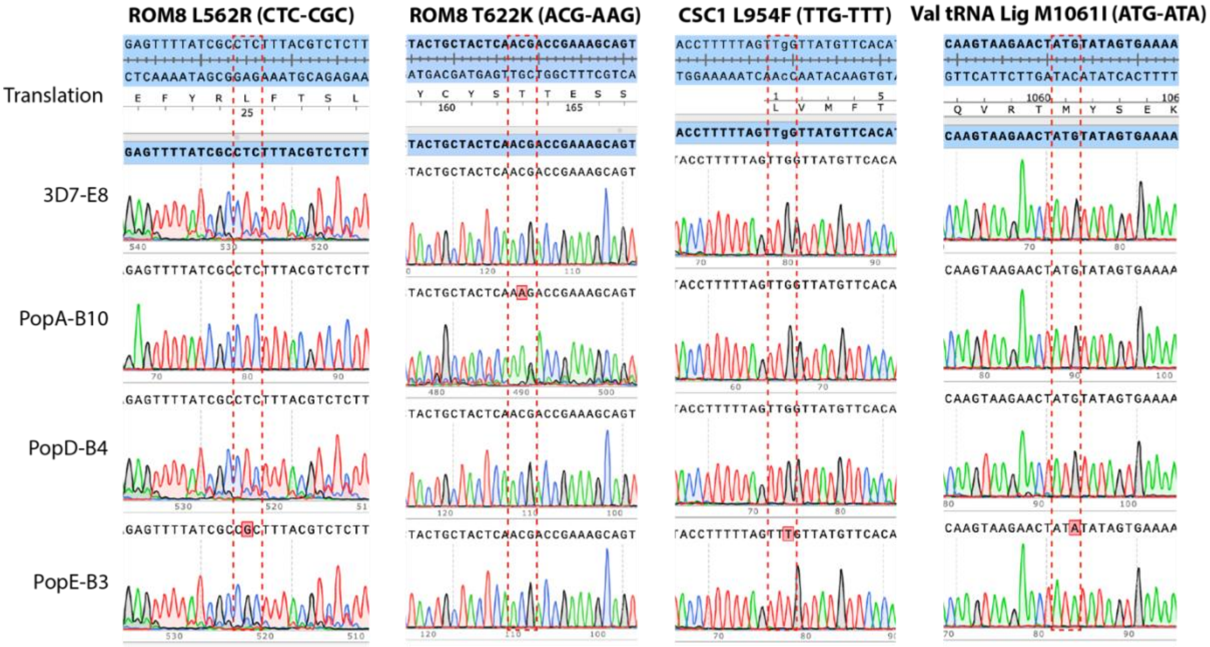


**Figure S3. Chromatograms of PCRs of genomic DNA from cloned MMV020512-resistant parasites from populations A, D and E confirming the relevant base changes in particular protein codons.** Chromatograms were generated in Snapgene V5.3.3.

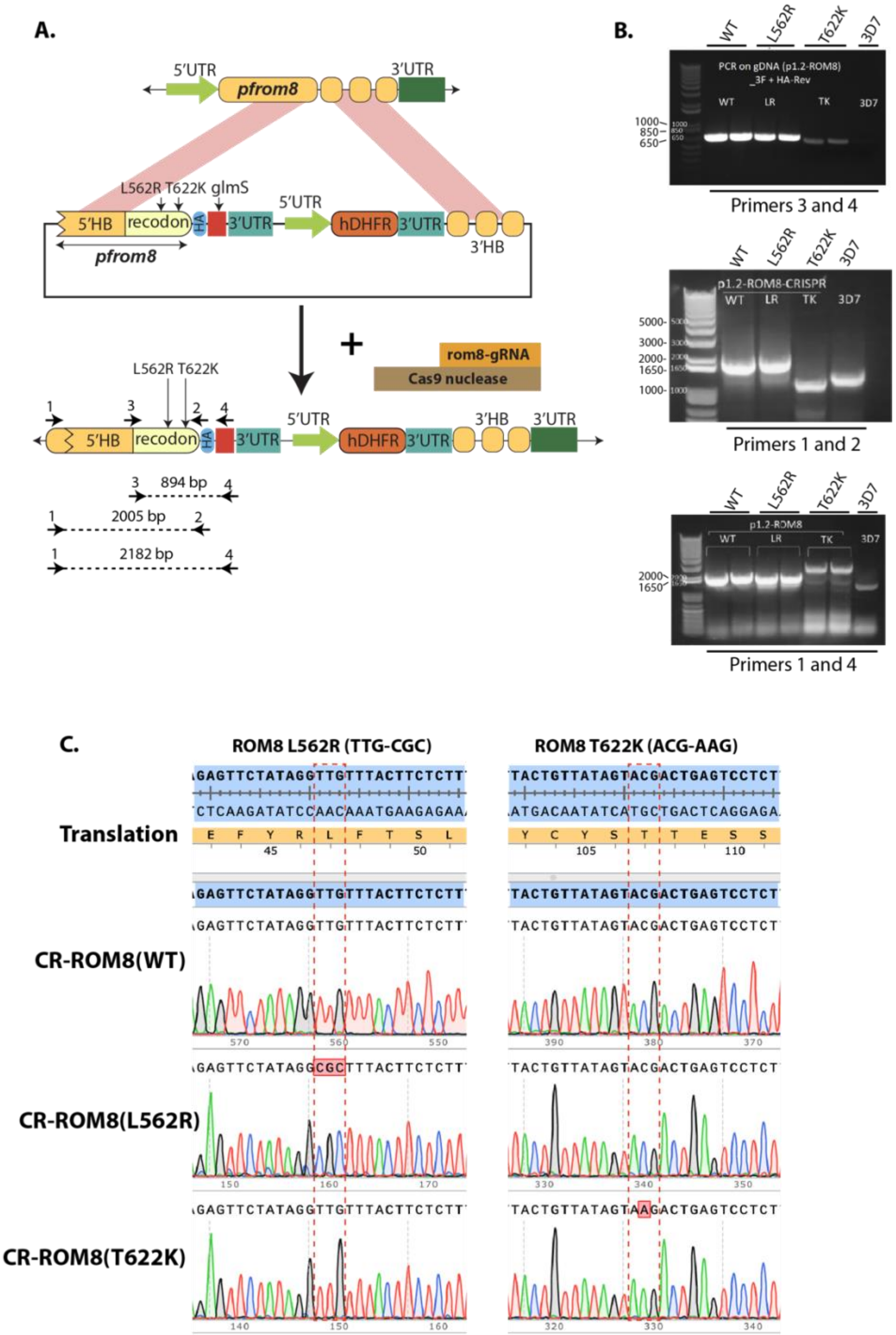


**Figure S4. Genetic modification of the rom8 locus. A)** Maps of *rom8* locus (top), the donor plasmid with L562R and T622K (middle) and the *rom8* locus modified with donor plasmid (bottom). Primer binding sites for diagnostic PCRs shown below. **B)** Diagnostic PCRs produced correctly sized products from genomic DNA of CR-ROM8(WT) and CR-ROM8(L562R) parasites but not for CR-ROM8(T622K) and 3D7 parasites. **C)** Chromatograms of sequencing reactions derived from PCR (IntF and HAglmS_R) of transfected parasites indicated that although the correct mutations were present, integration into the *rom8* locus was only correct for CR-ROM8(WT) and CR-ROM8(L562). Chromatograms produced in Snapgene V5.3.

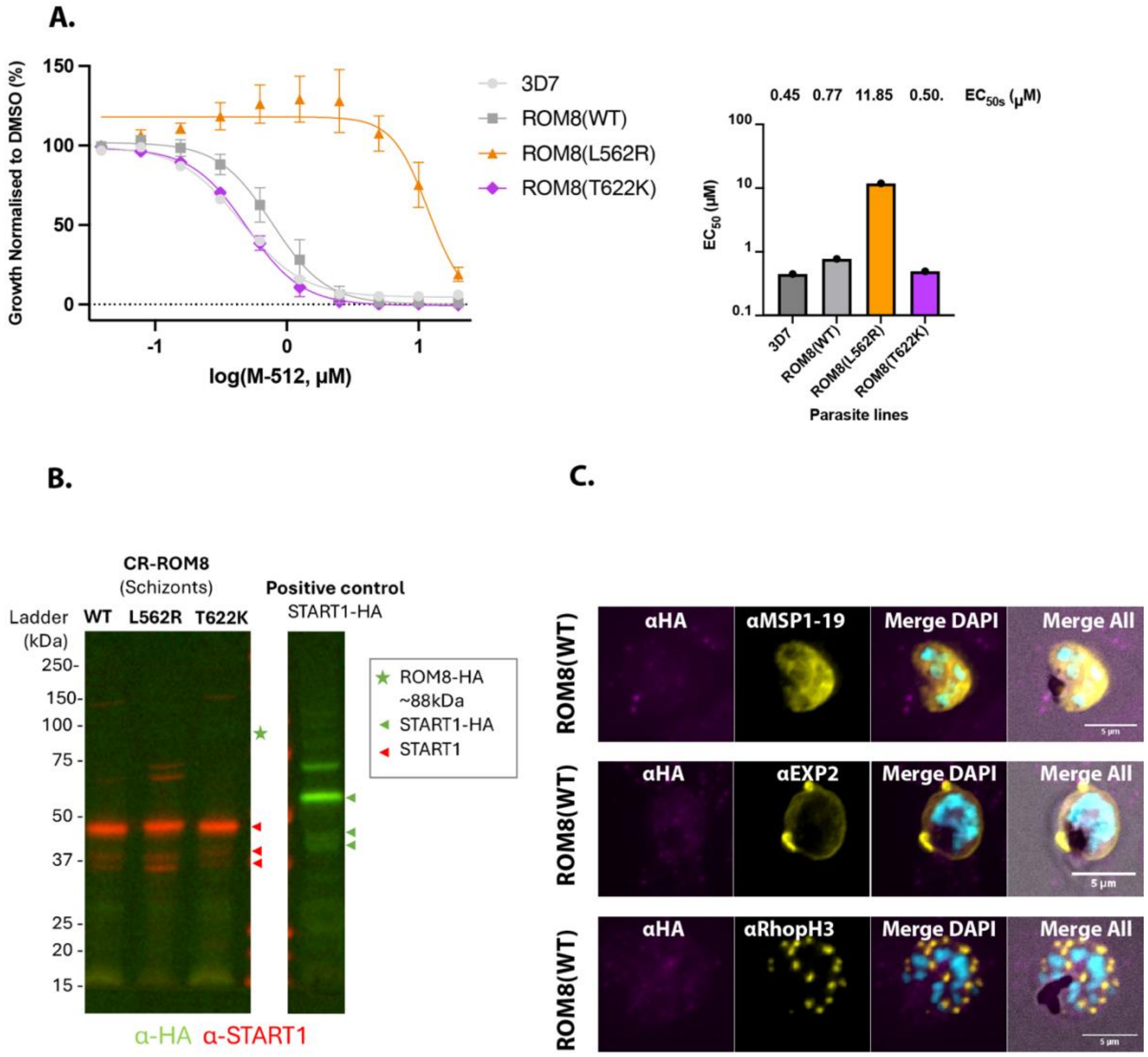


**Figure S5. Insertion of the L562R mutation into the *rom8* locus reproduces resistance to M-512 observed with an identical naturally acquired *rom8* resistance mutation. A)** The LDH activity of parasites treated with M-512 for 72-hours indicates that insertion of the L562R mutation into the *rom8* locus produces more resistance to M-512 compared 3D7 and CR-ROM(WT) and the incorrectly tagged CR-ROM8(T622K) parasites. Mean $EC_{50}$ values of technical triplicates of one biological replicate (Right). **B)** Western blots of CR-ROM8(WT) and *Pf*START1-HA parasites indicate HA-tag appended to the end of the CR-ROM8(WT) protein was not detected (green star) in contrast to the *Pf*START1-HA protein (green arrowheads). The loading control for CR-ROM8(WT) was the native *Pf*START1 protein detected with a protein specific antibody (red arrowheads). **C)** Immunofluorescence microscopy of CR-ROM8(WT) parasites did not detect a HA signal compared to antibodies to endogenous MSP1-19, EXP2 and RhopH3 proteins.

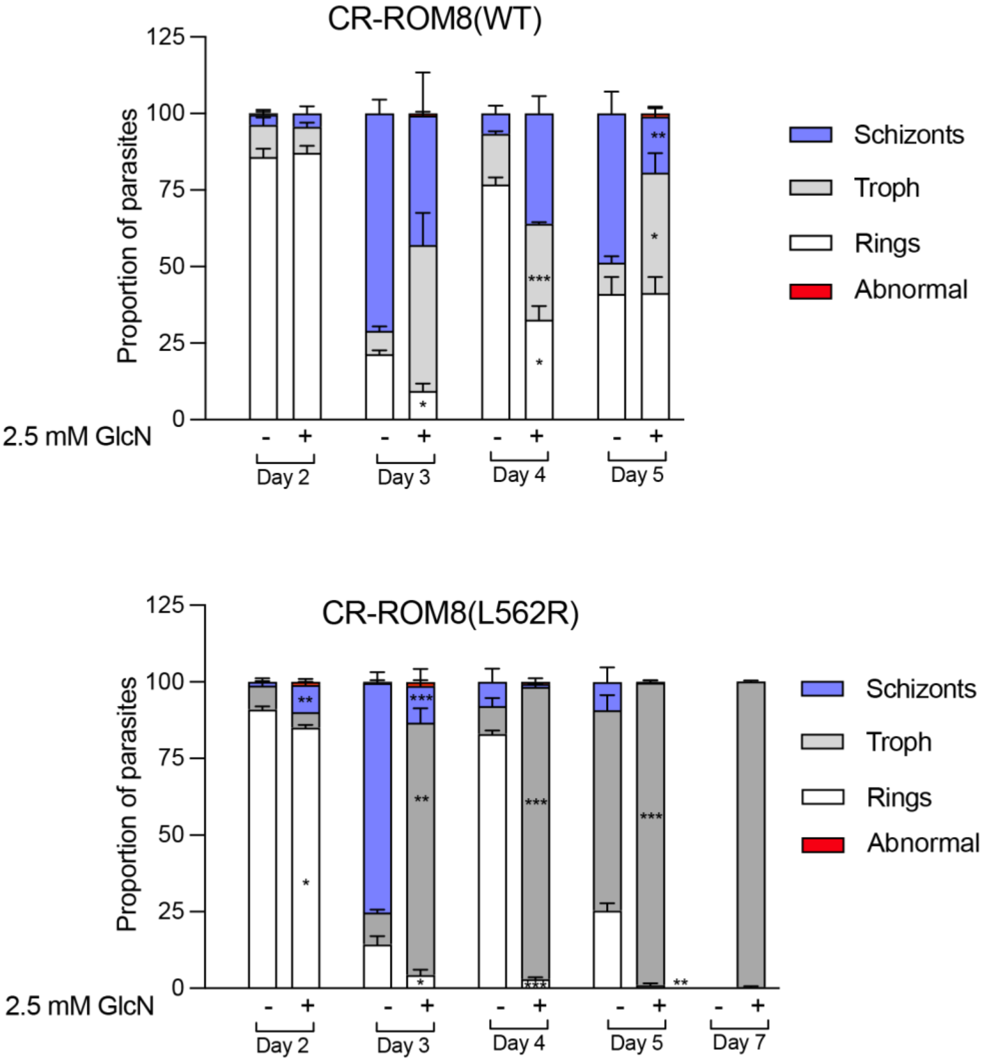


**Figure S6. Knockdown of ROM8 slows or arrests parasite growth three days after induction of the knockdown.** 2.5 mM glucosamine (GlcN) was added to ring-stage CR-ROM8(WT) and CR-ROM8(L562R) parasites on Day 0. Thin Giemsa-stained blood smears were made every two days and the numbers of rings, trophozoites, schizonts and abnormal parasites were counted and the proportions of each are shown. Compared to parasites not treated with GlcN, GlcN-treated CR-ROM8(WT) appeared to be growing more slowly by Day 3 with CR-ROM8(L562R) growth appearing to arrest by this time.

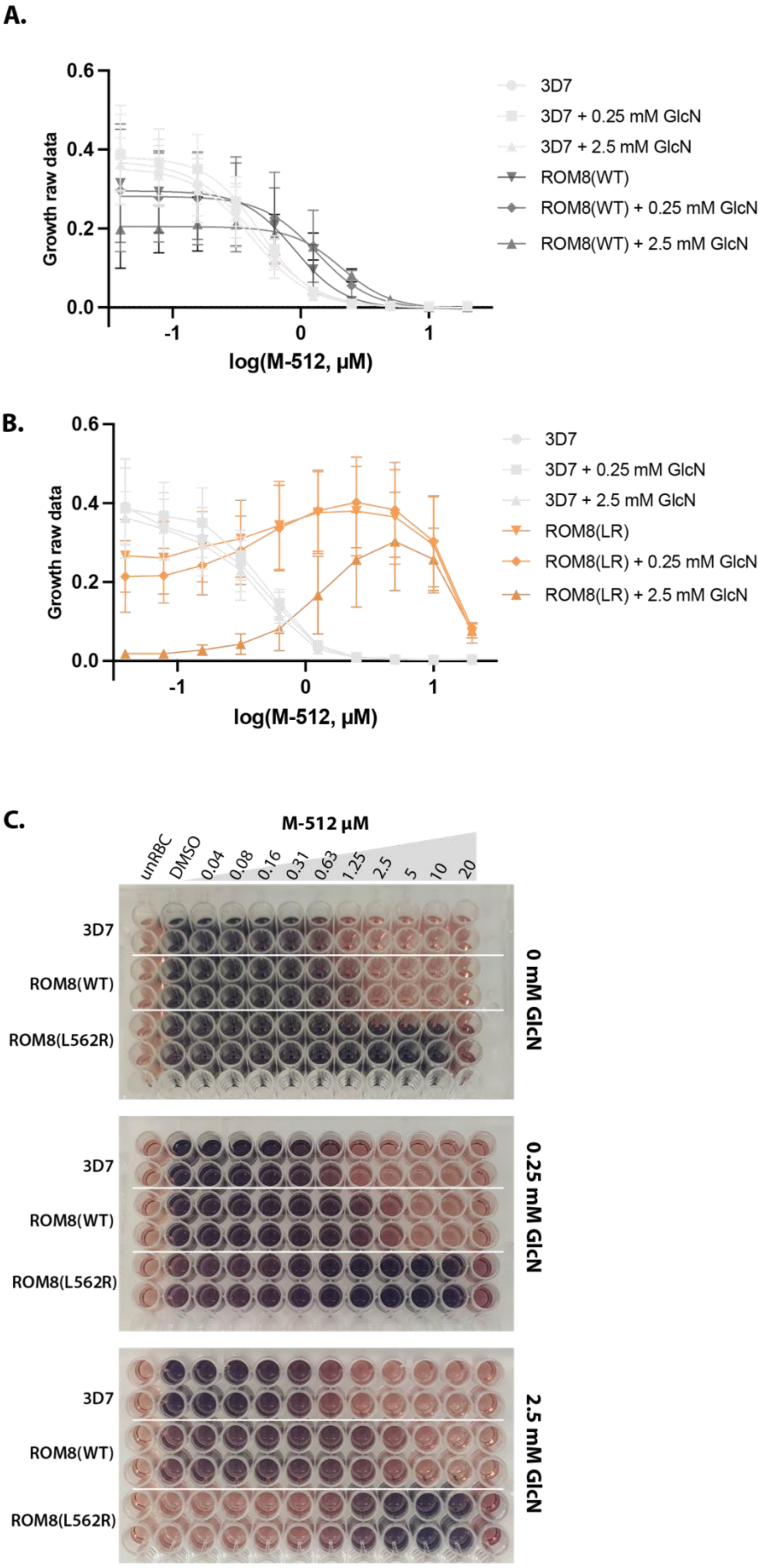

A.
0.6
0.4
0.2
0.0
Growth raw data
-1
0
1
log(M-512, µM)
3D7
3D7 + 0.25 mM GlcN
3D7 + 2.5 mM GlcN
ROM8(WT)
ROM8(WT) + 0.25 mM GlcN
ROM8(WT) + 2.5 mM GlcN
B.
0.6
0.4
0.2
0.0
Growth raw data
-1
0
1
log(M-512, µM)
3D7
3D7 + 0.25 mM GlcN
3D7 + 2.5 mM GlcN
ROM8(LR)
ROM8(LR) + 0.25 mM GlcN
ROM8(LR) + 2.5 mM GlcN
C.
M-512 µM
unRBC
DMSO
0.04
0.08
0.16
0.31
0.63
1.25
2.5
5
10
20
3D7
ROM8(WT)
ROM8(L562R)
0 mM GlcN
3D7
ROM8(WT)
ROM8(L562R)
0.25 mM GlcN
3D7
ROM8(WT)
ROM8(L562R)
2.5 mM GlcN

**Figure S7. M-512 overcomes the growth reduction due to knockdown of ROM8. A)** Growth inhibitory curves of 3D7 and CR-ROM8(WT) parasites treated with either 0, 0.25 or 2.5 mM GlcN and a dilution series of M-512 for 72 hours. The data were normalised and variable slope linear regression curves were fitted in PRISM V10. The graph is the same as that shown in Figure 4A but includes error bars indicating SD values. **B)** Growth inhibitory curves of 3D7 and CR-ROM8(L562R) parasites treated with either 0, 0.25 or 2.5 mM GlcN and a dilution series of M-512 for 72 hours. The data were normalised but linear regression curves could not be fitted for ROM8(LR) and data points were connected with lines. Mean values of 3 biological replicates in technical duplicate are indicated by data points and whisker indicate SD. **C)** Example plates of lactate dehydrogenase (LDH) activity of one of the biological replicates shown in A) and B). Increasing levels of parasite growth are indicated by a more intense pink to purple colour change. The data is graphically displayed in Figures 3 and 4.

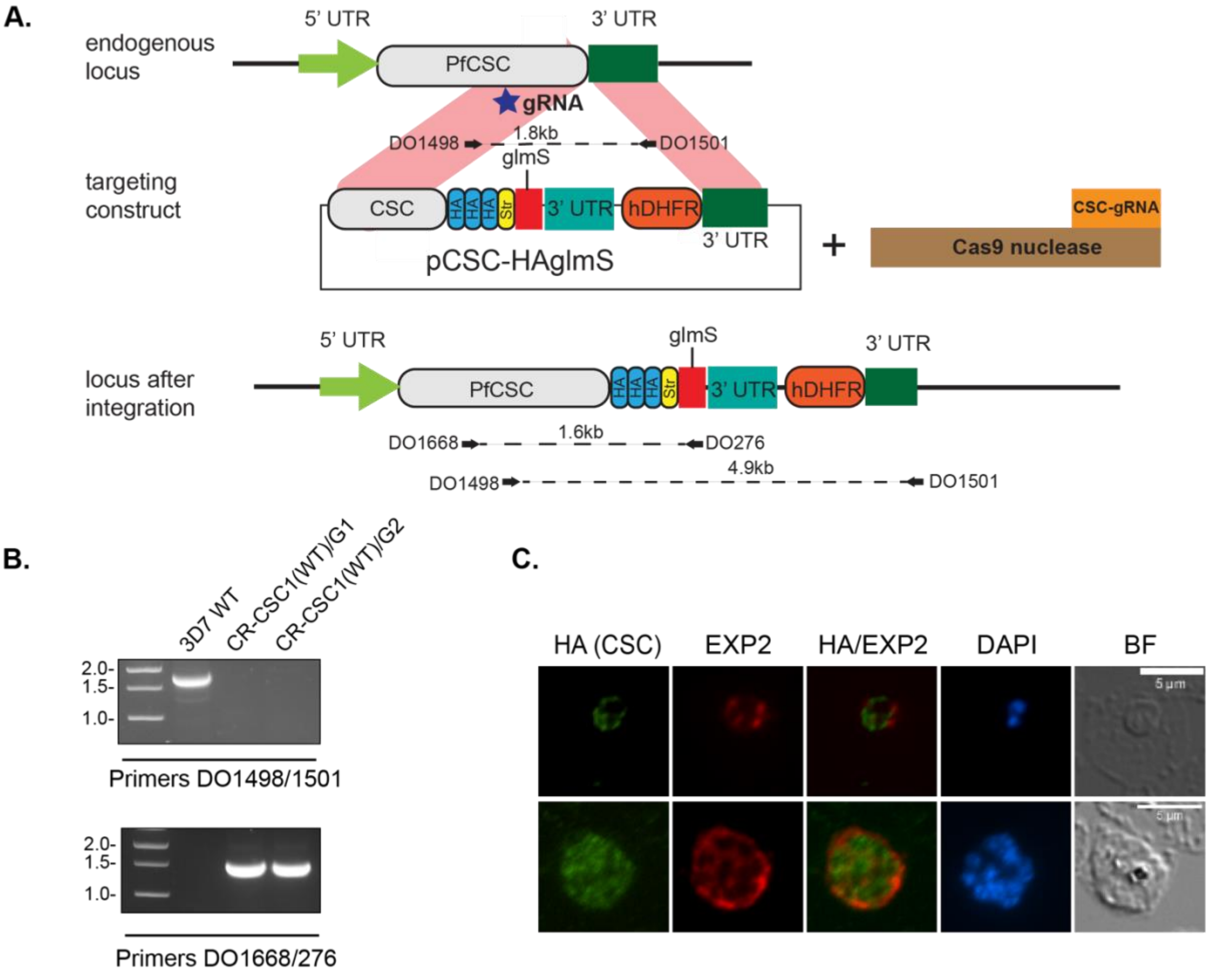


**Figure S8: Generation of PfCSC-HAglmS transgenic parasites. A)** Schematic of the *csc1* locus, and locus after double crossover recombination using CRISPR/Cas9. HA, haemagglutinin tag; Str, strep II tag. The plasmid also includes a *glmS* ribozyme with a heterologous untranslated region (UTR) and the selectable marker human dihydrofolate reductase (hDHFR). Arrows indicate oligonucleotides used in diagnostic PCRs as well as their expected sizes. **B)** Diagnostic PCR confirming integration of pCSC-HAglmS at the endogenous locus two transgenic lines in which two different gRNAs were used. (C) Immunofluorescence analysis of RBCs infected with CR-ROM8(WT) parasites using anti-HA antibodies to detect CSC1 and EXP2, which labels the PVM.

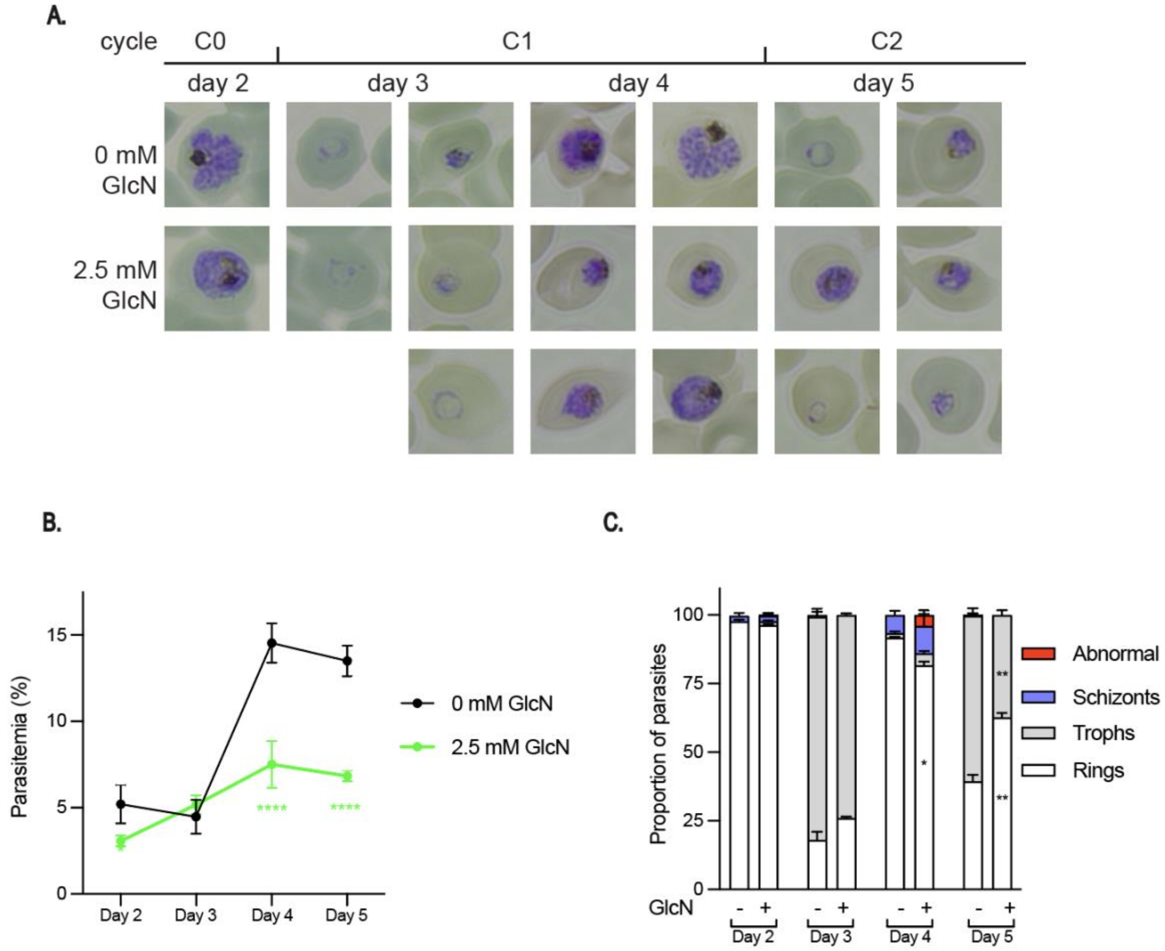


**Figure S9: Knockdown of CSC1 expression leads to perturbed parasite growth *in vitro*. A)** Representative Giemsa-stained parasite smears of CR-CSC1(WT) with a growth delay following reinvasion into cycle 2 (C2). In 2.5 mM GlcN, some parasites arrested at the trophozoite stage (middle row) while other parasite were able to progress to the next cycle (bottom row). **B)** Percentage parasitemia of CR-CSC1 +/- GlcN shows a growth defect by day 4 (end of C1). (n = 3 biological replicates, 500 cells counted per replicate). Shown is the mean ± standard deviation (n= 3). Statistical significance was analysed using 2-way ANOVA comparing parasite lines at each timepoint, followed by Sidak's multiple comparisons test (*** $p \leq 0.001$; ****$p \leq 0.0001$) **C)** Proportion of parasites at each stage +/- GlcN shows that depletion of CSC1 leads to delayed parasite development within cycle 2 (day 4; n = 3 biological replicates, 100 cells counted per replicate). Shown is the mean ± standard deviation (n= 3). Statistical significance was analysed unpaired t-tests at each timepoint (*** $p \leq 0.001$; ****$p \leq 0.0001$).
